# O-GlcNAc clusters attenuate BRD4 phase separation for transcription regulation

**DOI:** 10.1101/2025.10.20.683410

**Authors:** Guojian Shao, Qidong Deng, Tianyu Wang, Cuihua Li, Na Wang, Jun Liu, Yun Ge

## Abstract

Post-translational modifications (PTMs) regulate the liquid–liquid phase separation (LLPS) that organizes biomolecular condensates essential for transcription, yet for the monosaccharide O-GlcNAc, the general rules and their links to protein function remain undefined, with evidence largely limited to isolated cases rather than systematic analyses. Here we combine proteome-scale analyses with experiments to show that O-GlcNAc is a widespread LLPS modulator: glycosites are enriched in intrinsically disordered regions and form dense O-GlcNAc clusters. Using BRD4 as a model, we identified twelve clustered sites in its C-terminal IDR and showed that O-GlcNAc decreases condensate size while increasing fluidity in vitro and in cells. Removing O-GlcNAc strengthened BRD4 binding at active enhancers and promoted LLPS-mediated co-recruitment of transcriptional factors, including YTHDC1, leading to altered expression programs linked to the cell cycle and DNA repair. Our findings define O-GlcNAc clusters as regulators of condensate material properties and transcriptional outcomes, supporting a general paradigm in which PTMs fine-tune the molecular grammar of biomolecular condensates.

## Introduction

Biomolecular condensates, dynamic membraneless assemblies formed via liquid–liquid phase separation (LLPS), have emerged as a third layer of cellular compartmentalization, complementing membrane-bound organelles and classical protein complexes^1,2^. By concentrating factors in space and time, condensates govern essential biological processes, including transcription, signal transduction and stress adaptation^3–5^. For instance, in the nucleus, transcription factors, co-activators, and RNA polymerase II (Pol II) coalesce into micron-scale “transcriptional hubs”, whose material properties shape the magnitude and kinetics of gene expression^6–9^. Domain–ligand interactions and intrinsically disordered regions (IDRs) are central to the multivalent cooperativity that drives condensate formation^10^. The widespread occurrence of IDRs in phase-separating proteins underscores the importance of deciphering the molecular basis of IDR-mediated interactions^11^. While increasing biophysical studies have begun to define the “molecular grammar” encoded by amino acid composition and repeat motifs^12–14^, a comprehensive regulatory logic for how cells fine-tune IDR interactions—and consequently, condensate assembly, dynamics, and material properties—remains under exploration.

Notably, protein sequence features of IDRs also render them frequent targets of post-translational modifications (PTMs), which serve as “diacritical marks” that reprogram the primary molecular grammar with enhanced chemical diversity^15–17^. Like LLPS itself, PTMs are typically dynamic and reversible, allowing proteins to respond rapidly to environmental signals. PTMs modulate charge, hydrophobicity, and secondary structure of the primary sequence, thereby reshaping protein–protein and protein–RNA interactions^18–20^. Various PTMs—including phosphorylation, methylation, and poly(ADP-ribosyl)ation—have been extensively identified on IDRs, where they regulate LLPS by altering π-π stacking, cation-π, electrostatic and hydrophobic interactions^21–24^. For example, phosphorylation on transcriptional regulators can either promote or suppress condensate formation, modulate condensate partitioning and miscibility, and ultimately impact gene expression^22,25,26^. Moreover, phosphorylation patterns such as charge blockiness have been associated with LLPS regulation^21,27,28^. However, unlike phosphorylation, the modification patterns and regulatory principles of other IDR-targeting PTMs, especially the ubiquitous monosaccharide modification O-linked β-*N*-acetylglucosamine (O-GlcNAc), remain poorly defined.

O-GlcNAcylation raises of particular interests due to its physicochemical uniqueness. It involves the addition of a single, hydrophilic, and neutral glycan to serine (Ser) or threonine (Thr) residues on thousands of nucleocytoplasmic proteins^29^. This modification is catalyzed by a single pair of cycling enzymes: O-GlcNAc transferase (OGT) and O-GlcNAcase (OGA). O-GlcNAcylation integrates nutrient sensing and cellular stress and preferentially modifies IDRs^30^. It has been shown to reduce amyloid aggregation, increase the fluidity of SynGAP and YTHDF1/3 condensates in cells, and modulate the phase separation of EWSR1 in vitro^31–34^. While these individual studies suggest a regulatory role for O-GlcNAc in protein LLPS^35^, proteome-wide understandings of how O-GlcNAc patterning intersects with IDR-mediated LLPS and how this modulation affects downstream biology, remain lacking. In this study, we sought to decode the “O-GlcNAc code” in IDRs that governs LLPS behavior.

Here, we systematically profile curated O-GlcNAc sites across the human proteome and uncover a strong correlation between O-GlcNAc density and protein LLPS propensity. We identify a motif enriched in IDRs—comprising dense and consecutive proline (Pro), Ser, and Thr residues—which led to the discovery of O-GlcNAc clusters. These clusters are highly associated with LLPS-prone transcriptional regulators, with BRD4 ranking among the top candidates. As a key co-activator, BRD4 forms condensates at super-enhancers (SEs) in coordination with other factors to drive robust transcription^36^. We identify O-GlcNAc clusters in the C-terminal IDR of BRD4 and demonstrate that these clusters suppress BRD4 phase separation both *in vitro* and in cells. O-GlcNAc-induced changes in BRD4 condensates significantly influence its genomic occupancy and transcriptional output, primarily at SEs. Moreover, BRD4 condensate remodeling is accompanied by chromatin binding alterations in other LLPS-related transcriptional regulators, suggesting a coordinated regulatory network. Notably, O-GlcNAcylation of BRD4 impacts gene expression programs related to cell cycle progression, proliferation, and DNA damage response. Importantly, loss of O-GlcNAcylation enhances HeLa cell resilience to DNA damage and causes accelerated proliferation. Collectively, our study provides a systematic analysis of O-GlcNAc modification patterns in IDRs and uncovers a functional role for O-GlcNAc clusters in modulating protein condensates. Given the critical role of BRD4 as a coactivator, we further reveal a novel regulatory mechanism by which O-GlcNAc suppresses phase separation and fine-tunes transcriptional activity at enhancers, offering new insights into the PTM regulation of transcriptional condensates.

## Results

### O-GlcNAc clusters are highly correlated with protein phase separation ability

Mass spectrometry-based O-GlcNAc studies over the past decades have enabled the construction of a comprehensive human O-GlcNAcome, integrating available data on O-GlcNAc-modified proteins and sites^37^. To explore potential connections between O-GlcNAc modification and phase separation, we began by systematically analyzing the curated glycosite data in this database. Using the phase separation predictor SaPS and public phase-separating databases^38–41^ to estimate LLPS propensity, we categorized the human proteome into three groups: proteins with no predicted phase separation (PS) [Non-PS (predicted)], predicted PS [PS (predicted)], and experimentally reported PS [PS (reported)]. Compared with proteins lacking PS potential, those with either predicted or experimentally confirmed PS propensity exhibited significantly higher O-GlcNAc site levels (**Fig. 1a**). Further grouping proteins by the number of detected O-GlcNAc sites revealed that those with more glycosites tend to have higher SaPS scores (**Fig. 1b**), as well as a greater proportion of proteins with reported PS activity (**Extended Data** Fig. 1a). Subsequent analysis also revealed a strong positive correlation between O-GlcNAcylation density and the proportion of proteins with PS propensity (**Fig. 1c**). Given that proteins with PS propensity often contain IDRs critical for mediating multivalent interactions, we next analyzed glycosylation patterns across all IDR-containing proteins (DRPs). Proteins with more glycosites tended to have a greater proportion of DPRs, and compared to non-DRPs, DRPs displayed significantly higher O-GlcNAc site levels (**Fig. 1d, Extended Data** Fig. 1b). Moreover, DRPs with more glycosites exhibited a higher fraction of IDRs (**Fig. 1e**), and glycosite density positively correlated with the average IDR fraction in DRPs (**Fig. 1f**). These findings together suggest that increased O-GlcNAc sites are associated with a higher degree of intrinsic disorder and phase-separation potential.

**Fig. 1.**
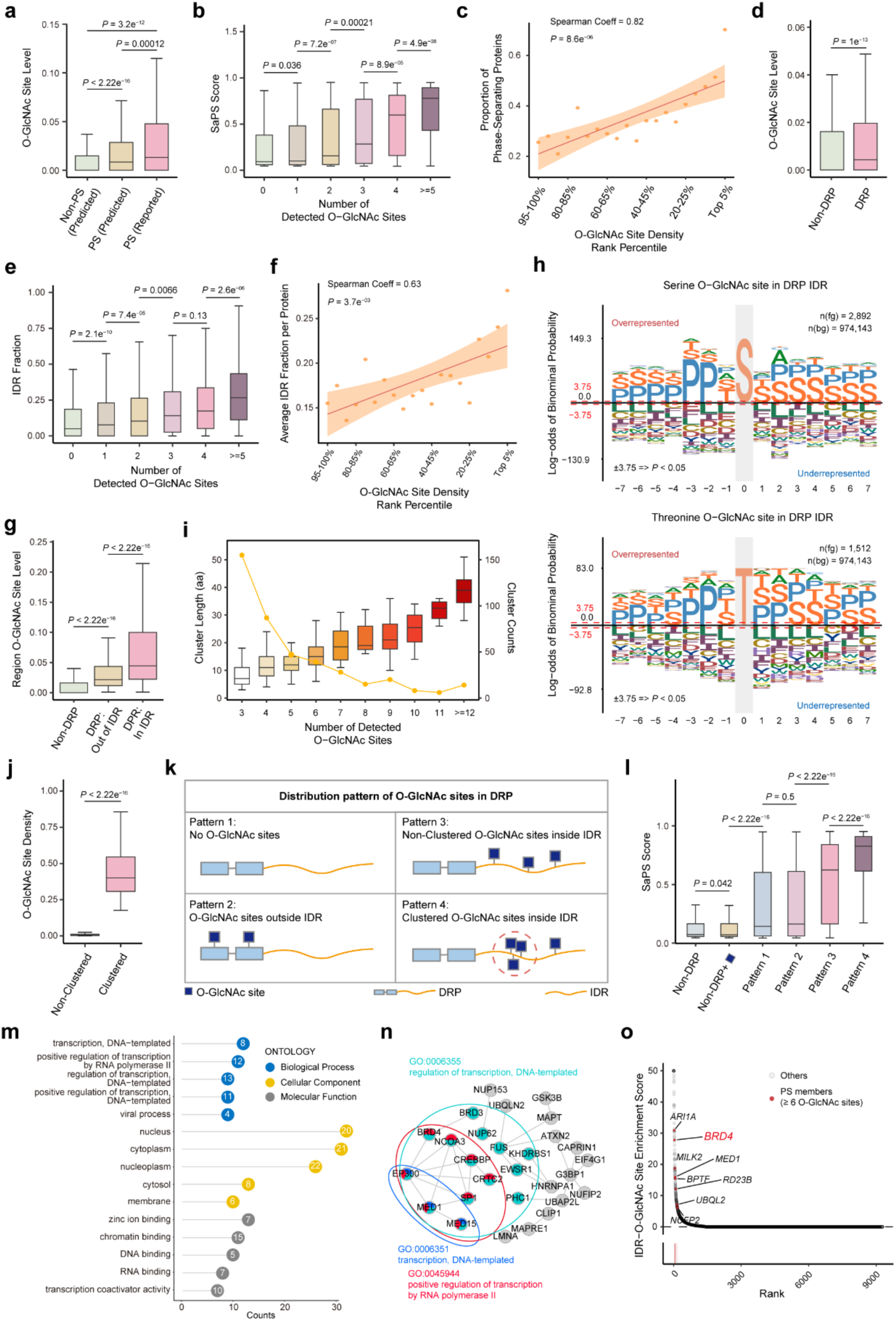
O-GlcNAc clusters are enriched in phase-separating proteins. **a.** Comparison of the O-GlcNAc site level among predicted Non-PS (predicted), PS (predicted) and PS (reported). **b.** Box plot depicting SaPS scores across protein groups containing different numbers of O-GlcNAc sites. **c.** The correlation between the rank percentile of O-GlcNAc site density and the proportion of phase-separating proteins. **d.** Box plot depicting the comparison of O-GlcNAc site level between IDR-containing proteins (DRP) and non-DRP. IDR, intrinsically disordered region. **e.** Box plot depicting the IDR fraction across protein groups containing different numbers of O-GlcNAc sites. **f.** The correlation between the rank percentile of O-GlcNAc site density and the average IDR fraction per protein. **g.** Box plot depicting comparison of region O-GlcNAc site levels among non-DRP, DRPs with glycosites outside and inside IDRs. **h.** O-GlcNAcylation consensus sequences of glycosites in IDRs of DRPs. O-GlcNAc sites are centered within a 15-amino acid window. **i.** Curve graph (yellow) and box plot showing the counts and length of O-GlcNAc clusters containing different numbers of detected O-GlcNAc sites. **j.** Box plot depicting comparison of O-GlcNAc site density between non-clustered sites and clustered sites. **k.** Schematic depiction of four O-GlcNAc distribution patterns in DRPs. Red circle highlights clustered O-GlcNAc sites in IDR. **l.** Box plot illustrating comparison of the SaPS scores among non-DRPs with and without O-GlcNAc, as well as DRPs with four distribution patterns as shown in **k**. **m.** Gene ontology (GO) analysis of 45 experimentally validated phase-separating proteins containing O-GlcNAc clusters. GO terms associated with molecular function, cellular component, and biological process are highlighted in gray, yellow, and blue circles with -log_10_ *P* inside, respectively. **n.** Protein-protein interaction (PPI) network of proteins analyzed in **m**, with proteins associated with three GO terms highlighted in green, red and blue. **o.** Plot showing the IDR-O-GlcNAc site enrich score O-GlcNAcylated proteins, with phase-separating proteins containing at least six clustered O-GlcNAc sites highlighted. The Spearman’s rank correlation coefficient (Spearman coeff) and *P* value are shown in **c, f**. Two-sided Wilcoxon rank-sum tests were used for statistical analyses in **a, b, d, e, g, j, l**, where boxes represent the 25^th^–75^th^ percentile with line at the median.

Furthermore, we divided DRPs into two groups based on whether their O-GlcNAc sites were located within IDRs or outside them. DRPs with glycosites in IDRs exhibited significantly higher O-GlcNAc site levels compared to those with sites outside IDRs (**Fig. 1g**). This prompted us to explore the sequence context of O-GlcNAc sites within DRPs. Although O-GlcNAc sites are known to lack a strict consensus motif (**Extended Data** Fig. 1c), alignment of glycosylated sequences in DRP IDRs revealed a distinct pattern enriched in Pro and Ser, forming dense and repetitive motifs surrounding the modified Ser or Thr residues (**Fig. 1h**). In contrast, glycosites outside IDRs or in non-DRPs displayed more diverse and less Pro/Ser-rich sequence contexts (**Extended Data** Fig. 1c). These Pro- and Ser/Thr-rich motifs suggest a tendency toward multisite O-GlcNAcylation within flexible, repetitive regions. Analyzing inter-site distances further confirmed that glycosites in DRP IDRs are more closely spaced than those in other regions, indicating clustered sites (**Extended Data** Fig. 1d). Therefore, we searched for O-GlcNAc clusters, defined as regions containing at least three O-GlcNAc sites within a 10–amino acid window. In total, we identified 419 O-GlcNAc clusters in IDRs across 308 DRPs, comprising 2,141 clustered glycosites—accounting for 47.3% of all glycosites in DRP IDRs (**Extended Data** Fig. 1e). These clusters ranged from 3 to 56 amino acids in length, with each cluster containing 3 to 24 glycosites (**Fig. 1i**). Compared to non-clustered sites, clustered glycosites showed significantly higher local O-GlcNAc site density (**Fig. 1j**).

To determine whether these clusters are functionally associated with phase separation, we further classified DRPs into four categories: (1) no glycosites, (2) glycosites outside IDRs, (3) non-clustered glycosites within IDRs, and (4) clustered glycosites within IDRs (**Fig. 1k**). SaPS scores were substantially higher in DRPs with glycosites in IDRs, and highest in those with O-GlcNAc clusters (**Fig. 1l**). A similar trend was observed in the proportion of proteins with reported PS abilities (**Extended Data** Fig. 1f), highlighting a strong association between O-GlcNAc clusters in IDRs and PS propensity.

Encouraged by these results, we further explored the functions of DRPs containing O-GlcNAc clusters. Among the 308 DRPs containing O-GlcNAc clusters, 45 were experimentally validated PS proteins. Gene Ontology (GO) analysis of these 45 proteins revealed enrichment in transcription regulation and chromatin binding, consistent with dense interaction networks in nuclear transcription (**Fig.1m, n**). Similar GO enrichment patterns were observed among all 308 proteins with O-GlcNAc clusters (**Extended Data Fig.1g, h**), collectively underscoring the potential functional relevance of these O-GlcNAc clusters in transcriptional regulation. We then developed an IDR-O-GlcNAc site enrichment score to integrate IDR length and O-GlcNAc site levels (**Fig.1o**). Notably, among DRPs with at least six clustered O-GlcNAc sites and known PS ability, most were transcription-related proteins, with Bromodomain-containing protein 4 (BRD4) ranked second. Altogether, our systematic analysis of O-GlcNAcylation patterns reveals that O-GlcNAc clusters are highly enriched within IDRs and are strongly associated with increased PS propensity. Importantly, proteins bearing O-GlcNAc clusters in IDRs are preferentially involved in transcriptional regulation, suggesting that O-GlcNAc clusters may play a key role in modulating condensate formation and function, particularly in RNA Pol II-mediated transcription.

### BRD4 C-terminal IDR is primarily modified with O-GlcNAc clusters

Given BRD4’s enrichment of multiple O-GlcNAc clusters and its known PS capability, we focused on BRD4 for further investigation. BRD4 plays critical roles in regulating gene expression^42^. However, beyond the information integrated in the human O-GlcNAc database, comprehensive biochemical validation of BRD4 O-GlcNAcylation remains limited. We first examined O-GlcNAc modification on endogenous BRD4 in HeLa cells. Endogenous BRD4 showed moderate O-GlcNAcylation, which increased nearly two-fold upon treatment with the OGA inhibitor Thiamet-G (TMG) and was slightly reduced by the OGT inhibitor OSMI-4 (**Fig. 2a, b**). These results indicate that BRD4 is dynamically O-GlcNAcylated under the regulation of OGT and OGA, with total protein levels remaining constant across conditions. BRD4 contains tandem N-terminal bromodomains (BD1 and BD2) that read acetyl-lysine marks, a central extra-terminal (ET) domain, and an extended C-terminal IDR. To identify the region responsible for interacting with OGT, we generated two HA-tagged BRD4 truncations: BRD4-NT (amino acids 1–719), which includes most of the structured domains, and BRD4-IDR (amino acids 674–1,351), encompassing the entire C-terminal IDR (**Fig. 2c**). Immunoprecipitation experiments in HEK293T cells revealed that BRD4-IDR, but not BRD4-NT, interacted with overexpressed His-tagged OGT (OGT-His) (**Fig. 2d**), echoing previous reports that OGT preferentially interacts with extended IDRs^30^. We next examined O-GlcNAcylation on these truncations by co-expressing them with either wild-type OGT or a catalytically inactive mutant [OGT(K852A)]. Only BRD4-IDR exhibited robust O-GlcNAcylation, while BRD4-NT showed no detectable modification (**Fig. 2e and Extended Data** Fig. 2a). This was further confirmed using a chemoenzymatic labeling and enrichment assay (**Extended Data** Fig. 2b, c). Additionally, we purified EGFP-fused BRD4-IDR (EGFP-IDR) and BRD4-NT (EGFP-NT) from *E. coli* as non-glycosylated controls, and performed *in vitro* O-GlcNAcylation using purified OGT and UDP-GlcNAc. Consistent with cellular results, EGFP-IDR was efficiently glycosylated, while EGFP-NT remained unmodified (**Fig. 2f, Extended Data** Fig. 2d). Together, these results indicate that OGT predominantly interacts with and modifies BRD4 through BRD4 C-terminal IDR.

**Fig. 2.**
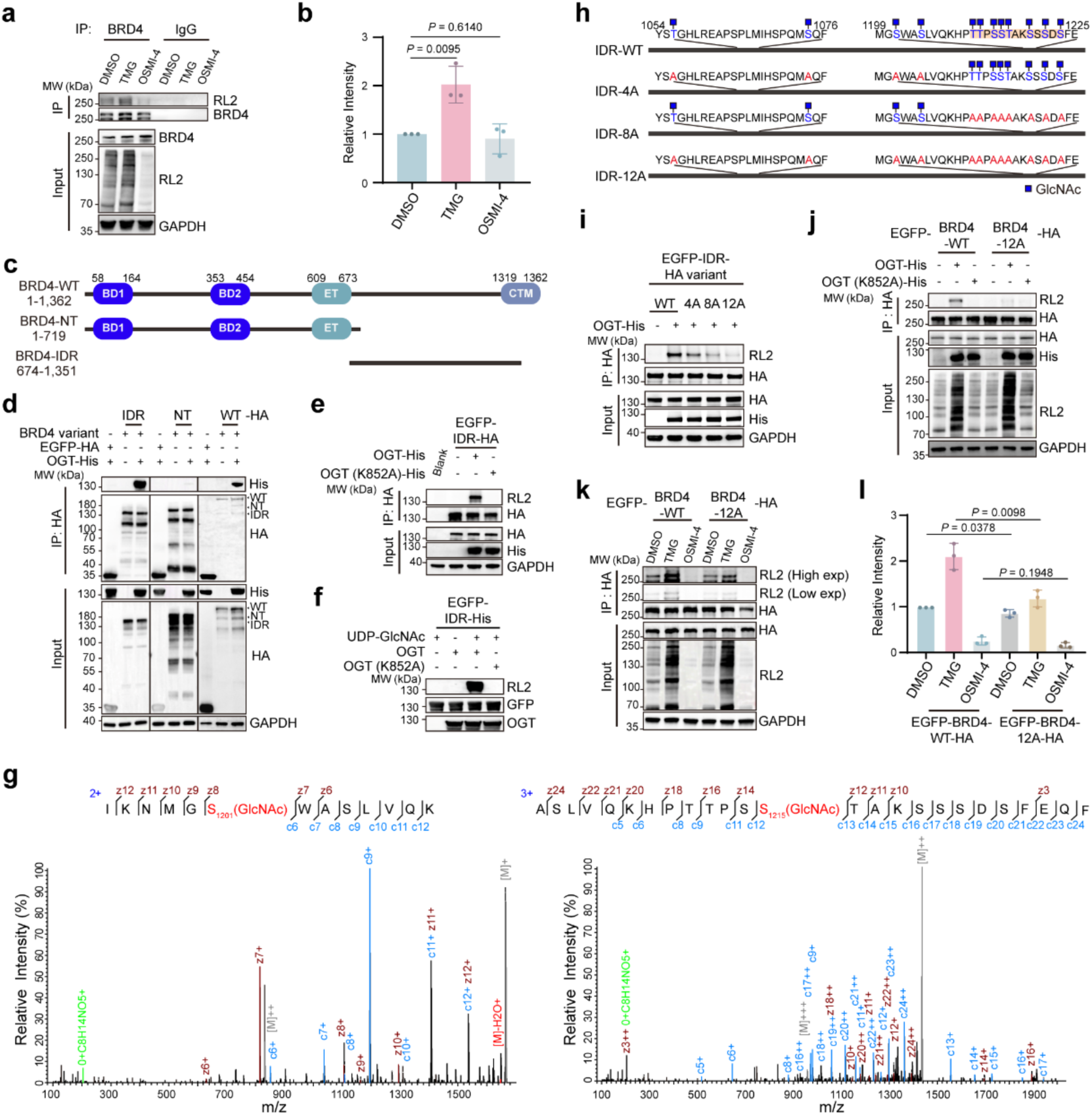
BRD4 C-terminal IDR interacts with OGT and modified with clustered O-GlcNAc. a, b. Immunoblots showing O-GlcNAc levels on endogenous BRD4 under Thiamet-G (TMG) or OSMI-4 treatments in HeLa cells. 20 μM TMG or OSMI-4 were added for 48 h. **c.** Schematic depiction of the domain architectures of BRD4-WT (1–1362), BRD4-NT (1–719), and BRD4-IDR (674–1351). BD1/2, Bromodomain 1/2; ET, extra terminal domain; CTM, C-terminal motif; WT, wild type; NT, N-terminus. **d.** Immunoblots showing interactions between OGT and BRD4 variants using the HA tag immunoprecipitation (IP) assay in HEK293T cells **e.** Immunoblots showing O-GlcNAc levels on HA and EGFP-tagged BRD4-IDR (EGFP-IDR-HA) when co-expressed with OGT or OGT (K852A) in HEK293T cells. **f.** Immunoblots showing O-GlcNAc level on purified EGFP-fused BRD4-IDR-His (EGFP-IDR-His) after *in vitro* O-GlcNAcylation assay. **g.** Representative mass spectra showing peptides on BRD4-IDR with O-GlcNAc modifications on S1201 (left) or S1215 (right). The c, z ions were annotated. **h.** Schematic depiction showing twelve O-GlcNAc sites and corresponding IDR variants with four (IDR-4A), eight (IDR-8A) or all twelve (IDR-12A) glycosites mutated to alanine. **i.** Immunoblots showing O-GlcNAc levels on EGFP-IDR-HA and indicated variants when co-expressed with OGT-His in HEK293T cells. **j.** Immunoblots showing O-GlcNAc levels on EGFP and HA-tagged full-length BRD4-WT (EGFP-BRD4-WT) and its mutant BRD4-12A (EGFP-BRD4-12A) when co-expressed with OGT-His in HEK293T cells. **k, l.** Immunoblots and quantification showing O-GlcNAc levels on EGFP-BRD4-WT and EGFP-BRD4-12A with TMG or OSMI-4 treatment for 48 h in HEK293T cells. RL2, anti-O-GlcNAc antibody (RL2). Exp, exposure. Data in **d–f, i, j** represent at least two biological replicates. Data are shown as the mean ± s. d. from n = 3 independent experiments. Two-tailed t-tests were used in statistical analyses.

Given the limited glycosite information in prior O-GlcNAc proteomics studies, we sought to comprehensively map potential glycosites on the BRD4-IDR. To maximize site coverage, EGFP-IDR purified from *E. coli* was subjected to *in vitro* O-GlcNAcylation followed by liquid chromatography with tandem mass spectrometry (LC-MS/MS) analysis. Ten putative glycosites on BRD4-IDR were identified (**Fig. 2g and Extended Data** Fig. 3), with seven located within a single chymotrypsin-digested peptide spanning A1203 to F1227. Considering possible misassignments of MS analyses and compensatory glycosylation, we included two additional adjacent glycosites (T1211 and S1223) previously reported in the O-GlcNAc database, bringing the total to 12 candidate glycosites for validation. Notably, the region spanning P1210 to S1223 aligns well with the Pro- and Ser-rich consensus motif identified earlier (**Fig. 1h**), and eight glycosites within this segment form a dense cluster (**Fig. 2h**).

To evaluate the contribution and fidelity of individual sites, we generated glycosylation-resistant BRD4-IDR mutants by substituting sites with alanine: 4A (first four sites), 8A (last eight sites), and 12A (all twelve sites) (**Fig. 2h**). Co-expression with active OGT in HEK293T cells showed a progressive decrease in O-GlcNAcylation with increasing numbers of mutated sites, with the 12A mutant showing nearly complete loss of modification (**Fig. 2i**). Consistent with this, LC-MS/MS analysis of *in vitro* glycosylated BRD4-IDR-12A failed to detect additional glycosylation sites (data not shown). Notably, reverting individual alanine mutations back to the original residues failed to restore O-GlcNAcylation even in the presence of OGT overexpression (**Extended Data** Fig. 2e), indicating that no single site dominates; instead, the full set of twelve sites cooperatively contributes to O-GlcNAcylation. Likewise, introducing the 12A mutations into full-length BRD4 (BRD4-12A) resulted in substantially reduced O-GlcNAcylation compared to wild-type BRD4 (BRD4-WT) in the presence of active OGT (**Fig. 2j**). BRD4-12A also showed diminished responsiveness to TMG treatment, reinforcing the notion that the C-terminal IDR is the primary site of dynamic O-GlcNAc modification (**Fig. 2k, l**). Despite abrogating IDR glycosylation, low levels of residual O-GlcNAc were detected in BRD4-12A, likely due to modification at other regions such as BRD4-NT, potentially mediated by IDR-facilitated interactions with OGT. Together, these results confirm the existence of clustered O-GlcNAc sites in the C-terminal IDR of BRD4 and demonstrate their dynamic regulation. The absence of dominant glycosites suggests a cooperative, compensatory mechanism, wherein moderate to low stoichiometry at individual residues collectively maintains a stable O-GlcNAc level within the cluster.

### O-GlcNAcylation suppresses BRD4 phase separation *in vitro*

BRD4 was recently shown to form phase-separated coactivator condensates at SEs together with Mediator, thereby facilitating RNA Pol II–mediated transcription^36^. Given the strong correlation between O-GlcNAc clusters and SaPS scores (**Fig. 1l**), we hypothesized that the O-GlcNAc clusters identified in the BRD4 C-terminal IDR modulate its phase separation behavior. To test this, we performed *in vitro* droplet formation assays. Consistent with previous reports, BRD4-IDR formed spherical droplets upon addition of 10% PEG8000. Droplet size decreased with increasing NaCl concentration (**Fig. 3a**) and increased with higher protein concentration (**Fig. 3c**). However, O-GlcNAcylation—achieved through incubation with UDP-GlcNAc and active OGT—significantly reduced droplet size while maintaining similar trends in response to NaCl and protein concentration (**Fig. 3b, d**). These results indicate that O-GlcNAcylation weakens BRD4-IDR phase separation, a finding corroborated by turbidity measurements (**Extended Data** Fig. 4a, b). Additionally, titration of unmodified BRD4-IDR with increasing proportions of O-GlcNAcylated BRD4-IDR resulted in progressively smaller droplets at constant total protein concentrations (**Fig. 3e**). This demonstrates that variation in O-GlcNAcylation stoichiometry provides a tunable mechanism for regulating droplet size, suggesting that cellular fluctuations in O-GlcNAc levels may dynamically modulate BRD4 phase behavior.

**Fig. 3.**
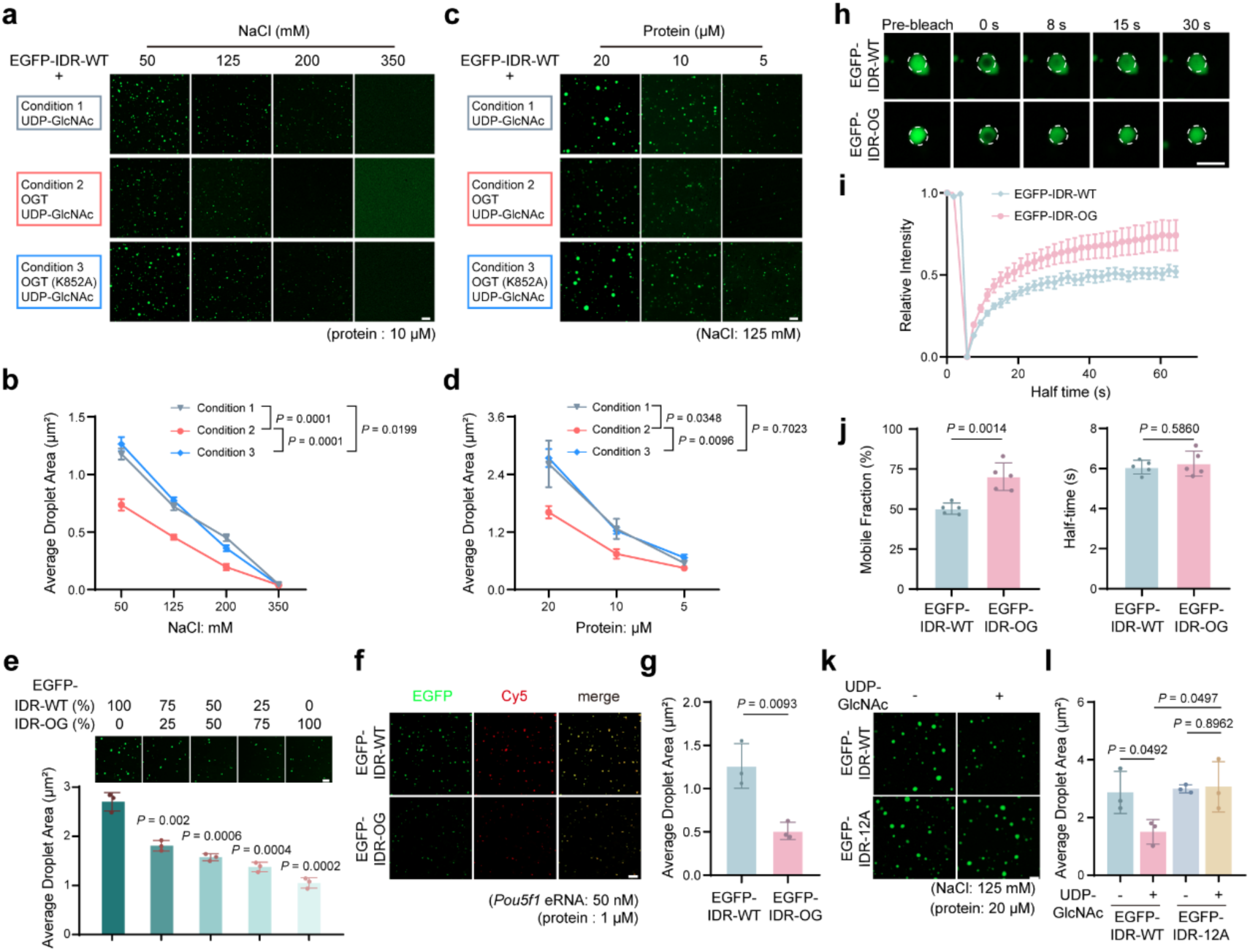
O-GlcNAcylation attenuates phase separation of EGFP-fused BRD4-IDR *in vitro*. **a, b.** Representative images (**a**) and quantification (**b**) of droplet formation of 10 μM EGFP-fused BRD4-IDR-His (EGFP-IDR-WT) under indicated *in vitro* O-GlcNAcylation conditions at varying NaCl concentrations. **c, d.** Representative images (**c**) and quantification (**d**) of droplet formation of EGFP-IDR-WT with varying protein concentrations under indicated *in vitro* O-GlcNAcylation conditions at 125 mM NaCl. **e.** Representative images (top) and quantification (bottom) of droplet formation with different O-GlcNAc-modified EGFP-IDR (EGFP-IDR-OG) molar percentages at a total of 10 μM protein concentration at 50 mM NaCl. **f, g.** Representative images (**f**) and quantification (**g**) of droplets formed by 50 nM *Pou5f1* enhancer RNA (*Pou5f1* eRNA, indicated in the Cy5 channel) mixed with 1 μM EGFP-IDR-WT or EGFP-IDR-OG. **h.** Fluorescence recovery after photobleaching (FRAP) analysis of EGFP-IDR-WT droplets (top) and EGFP-IDR-OG droplets (bottom) *in vitro*. **i.** FRAP curves for EGFP-IDR-WT and EGFP-IDR-OG. **j.** Histograms showing the mobile fraction (left) and recovery half-time (right) of droplets in **h**. Data in **h–j** are represented by the mean ± s.d. of n = 5 individual droplets from three biological independent samples. **k, l.** Representative images (**k**) and quantification (**l**) of droplets formed by 20 μM EGFP-IDR-WT and its 12A mutant (EGFP-IDR-12A) at 125 mM NaCl following *in vitro* O-GlcNAcylation in the presence or absence of UDP-GlcNAc. Data in **a–g, k, l** are represented by the mean ± s.d. from n = 3 biologically independent samples. Scale bar, 5 μm (**h**), 10 μm (**a, c, f, k**), and 20 μm (**e**). Statistical analyses were performed using two-way ANOVA tests in **b, d**, and two-tailed t-tests in **e, g, j, l**.

As a core component of transcriptional condensates, RNA is known to enhance BRD4-IDR phase separation at low concentrations but dissolve condensates at high concentrations^43^. In the presence of Cy5-labeled *Pou5f1* enhancer RNA, O-GlcNAcylated BRD4-IDR still exhibited this RNA-mediated biphasic response, although condensate size was attenuated overall (**Fig. 3f, g and Extended Data** Fig. 4c). This suggests that O-GlcNAcylation does not disrupt RNA-mediated feedback regulation of BRD4-IDR condensates. We speculate that this minimal interference may be due to the hydrophilic and neutral nature of O-GlcNAc, whereas RNA–protein co-condensation is primarily governed by electrostatic interactions.

We next assessed the material properties of BRD4-IDR condensates by fluorescence recovery after photobleaching (FRAP). O-GlcNAcylation led to increased fluorescence recovery after bleaching (**Fig. 3h–j**), indicating enhanced molecular mobility and more liquid-like dynamics within the condensates. Together, these findings show that while O-GlcNAcylation does not alter the general responsiveness of BRD4-IDR to salt, protein, or RNA concentrations, it attenuates condensate formation and increases condensate fluidity. This manifests as smaller droplet sizes and higher FRAP recovery rates *in vitro*, likely driven by the hydrophilicity, electronic neutrality, and steric hindrance introduced by O-GlcNAc clusters.

To further confirm that these effects were mediated by O-GlcNAcylation, we compared EGFP-fused BRD4-IDR-WT (EGFP-IDR-WT) to the glycosylation-resistant mutant BRD4-IDR-12A (EGFP-IDR-12A) in droplet assays. EGFP-IDR-12A formed droplets of similar size and fluidity to EGFP-IDR-WT under varying NaCl and protein concentrations (**Extended Data** Fig. 4d–h), indicating that the 12A mutations themselves did not perturb condensate formation. However, O-GlcNAcylation significantly reduced droplet size in EGFP-IDR-WT but had no effect on EGFP-IDR-12A (**Fig. 3k, l**), confirming that EGFP-IDR-12A is resistant to O-GlcNAc modification and that the observed effects are specific to O-GlcNAcylation of IDR.

### O-GlcNAcylation reduces phase-separated BRD4 in cells

We next investigated O-GlcNAc function in regulating BRD4 phase separation in cells. As expected, endogenous BRD4 formed liquid-like condensates in HeLa cells, which can be disrupted by 1,6-hexanediol (1,6-Hex), consistent with previous studies (**Fig. 4a**). Upon treatment with TMG to globally elevate O-GlcNAc levels, both the size and number of BRD4 nuclear puncta were significantly reduced (**Fig. 4a, b**). In contrast, treatment with the OGT inhibitor OSMI-4 had minimal effects, likely due to the inherently low baseline O-GlcNAcylation of endogenous BRD4 in HeLa cells. To minimize possible side effects from the global perturbations induced by the two inhibitors, we employed the nanobody-OGT for targeted protein O-GlcNAcylation^44^. Targeted O-GlcNAcylation of EGFP-BRD4-WT using the catalytically active nanobody–OGT fusion significantly reduced both the number and size of BRD4 nuclear puncta (**Fig. 4c, d, Extended Data** Fig. 5a), demonstrating that O-GlcNAcylation on BRD4 directly diminishes its condensate formation in cells.

**Fig. 4.**
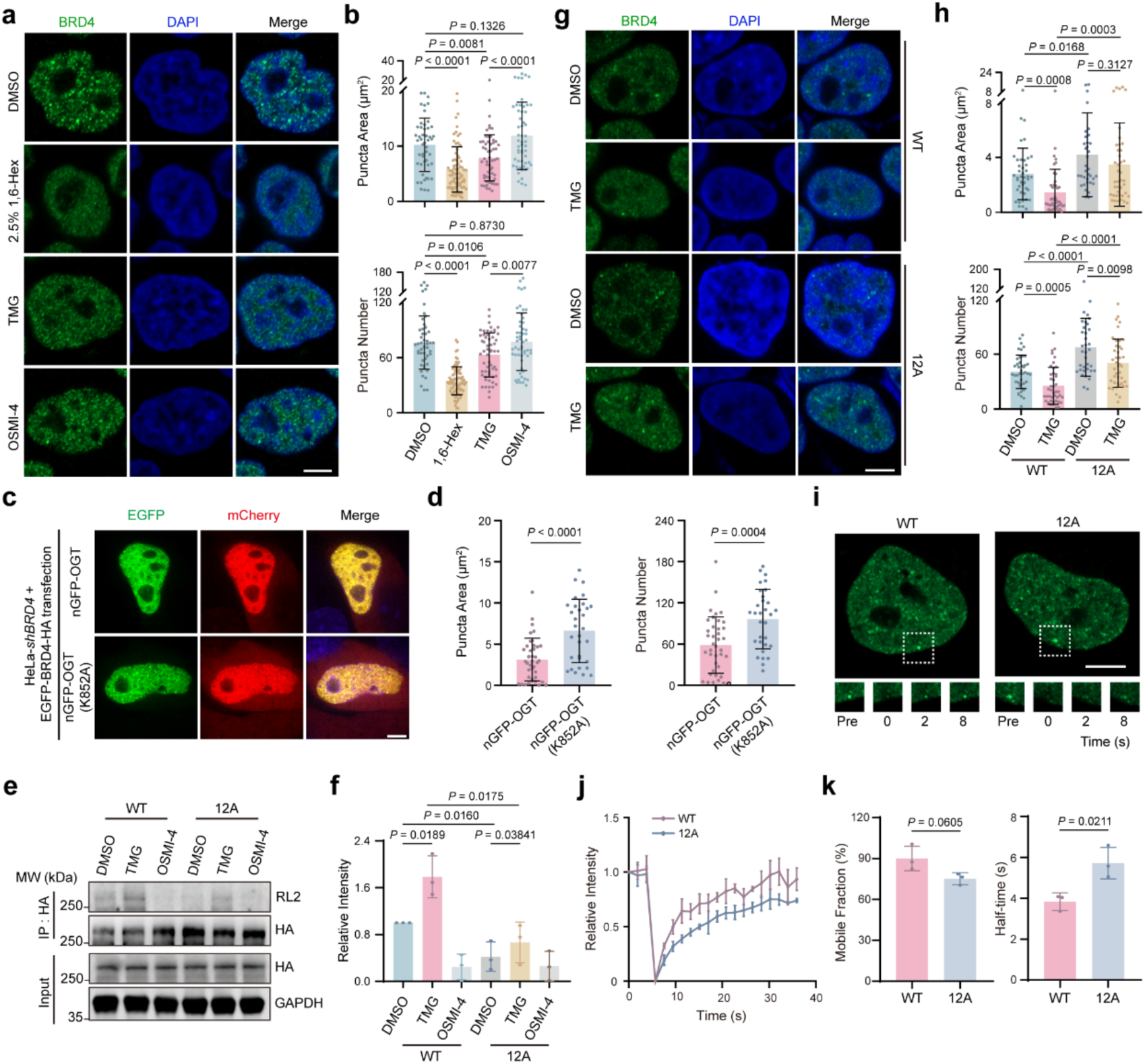
O-GlcNAcylation diminishes BRD4 condensation in HeLa cells. **a.** Representative immunofluorescence images showing endogenous BRD4 puncta in HeLa cells treated with 1,6-hexanediol (1,6-Hex), TMG, and OSMI-4. 2.5% 1,6-Hex was added for 10 min. **b.** Quantified puncta areas (top) and numbers (bottom) for samples in **a**. n = 49, 78, 57, and 54 cells per group, respectively, pooled from three independent replicates. **c.** Live-cell images showing the puncta formation of EGFP-BRD4-WT or EGFP-BRD4-12A when co-expressed with mCherry and anti-GFP nanobody-tagged OGT variants (nGFP-OGT) or its catalytically dead mutant [nGFP-OGT (K852A)] in *shBRD4* HeLa cells. **d.** Quantified puncta areas (left) and numbers (right) for samples in **c**. n = 39 and 31 cells per group, respectively, pooled from three independent replicates. **e, f.** Immunoblots showing O-GlcNAc levels of BRD4-WT (WT) and BRD4-12A (12A) stably expressed in *shBRD4* HeLa cells under inhibitors treatments. RL2, anti-O-GlcNAc antibody (RL2). **g.** Immunofluorescence images showing BRD4 WT and 12A cells with TMG treatment. **h.** Quantified puncta areas (top) and numbers (bottom) from samples in **g**. n = 42, 43, 35 and, 42 cells per group, respectively, pooled from three independent replicates. **i.** FRAP analysis of EGFP-fused WT or 12A condensates in rescue cell lines. Bottom, representative images showing a photobleached droplet at different time points. Pre, Pre-bleach. **j.** FRAP curves of EGFP-fused WT or 12A condensates from samples in **i**. **k.** Histograms showing the mobile fraction (left) and recovery half-time (right) of droplets in **i**. 20 μM TMG or OSMI-4 were added for 48 h in **a, e, g**. n = 3 biologically independent experiments in **f, j, k**. Scale bar, 5 μm (**a, c, g, i**). Data in **b, d, f, h, j, k** are represented by the mean ± s.d.. Two-tailed t-tests were used for statistical analyses.

To further investigate the functional role of O-GlcNAcylation, we established two stable HeLa cell lines in which endogenous BRD4 was knocked down and rescued with either EGFP-BRD4-WT or the glycosylation-deficient mutant EGFP-BRD4-12A, expressed at levels comparable to endogenous BRD4 (**Extended Data** Fig. 5b–d). As expected, EGFP-BRD4-WT exhibited significantly higher O-GlcNAc level than EGFP-BRD4-12A (**Fig. 4e, f**). TMG and OSMI-4 treatments respectively increased and decreased the O-GlcNAc level on EGFP-BRD4-WT, while EGFP-BRD4-12A showed minimal changes (**Fig. 4e, f**). Consistent with the glycosylation data, cells rescued with EGFP-BRD4-WT displayed significantly smaller and fewer condensates compared to those rescued with EGFP-BRD4-12A (**Fig. 4g, h**). Moreover, TMG treatment caused a substantial reduction in the size and number of EGFP-BRD4-WT puncta, but had little effect on EGFP-BRD4-12A. These observations were corroborated by transient transfection experiments in BRD4 knockdown HeLa cells expressing either EGFP-BRD4-WT or EGFP-BRD4-12A (**Extended Data** Fig. 5e–g). Overall, these results mirror our *in vitro* findings, where O-GlcNAcylation suppressed BRD4 droplet formation in a manner proportional to glycosylation levels.

Moreover, using FRAP analyses, we examined the liquid-like properties of BRD4 condensates in both cell lines. Consistent with in vitro results, EGFP-BRD4-WT exhibited a faster recovery rate and a moderately higher recovery ratio after photobleaching compared to the unmodified EGFP-BRD4-12A (**Fig. 4i–k**), indicating that O-GlcNAc on EGFP-BRD4-WT also promotes BRD4 liquidity in cells. Taken together, these in-cell findings, in agreement with *in vitro* evidence, demonstrate that dynamic O-GlcNAcylation at the BRD4 C-terminal IDR serves as a critical regulator of condensation properties.

### O-GlcNAcylation regulates genome-wide chromatin binding of BRD4

BRD4, a BET family member, binds acetylated histones and functions as a key transcriptional coactivator^45,46^. To determine whether O-GlcNAcylation regulates BRD4’s transcriptional activity by modulating its phase separation capacity, we first compared the chromatin binding ability of BRD4-WT and the O-GlcNAc-resistant BRD4-12A variant. Chromatin fractionation revealed increased BRD4-12A binding relative to BRD4-WT, with TMG treatment further reducing BRD4-WT occupancy (**Fig. 5a**), indicating that O-GlcNAcylation directly modulates BRD4’s chromatin occupancy. We next performed BRD4 chromatin-immunoprecipitation coupled to high-throughput sequencing (ChIP-seq) in HeLa cells stably expressing either BRD4-WT or BRD4-12A (**Extended Data** Fig. 6a). BRD4 peaks were predominantly localized to active enhancers (**Fig. 5b**). Compared to BRD4-WT, BRD4-12A exhibited significantly elevated ChIP-seq signal at active enhancers and promoters, but not at primed enhancers (**Fig. 5c**), suggesting that O-GlcNAcylation limits BRD4 occupancy at transcriptionally active regulatory elements. Furthermore, we identified 14,767 BRD4-bound typical enhancers (TEs), marked by H3K27ac and H3K4me1, and 484 BRD4-defined SEs using the ROSE algorithm (**Extended Data** Fig. 6b). BRD4-12A showed stronger occupancy at 10,448 TEs and across SEs and promoters compared to BRD4-WT (**Fig. 5d, e, Extended Data** Fig. 6c), reinforcing the role of O-GlcNAcylation in reducing BRD4 chromatin binding, particularly at enhancers.

**Fig. 5.**
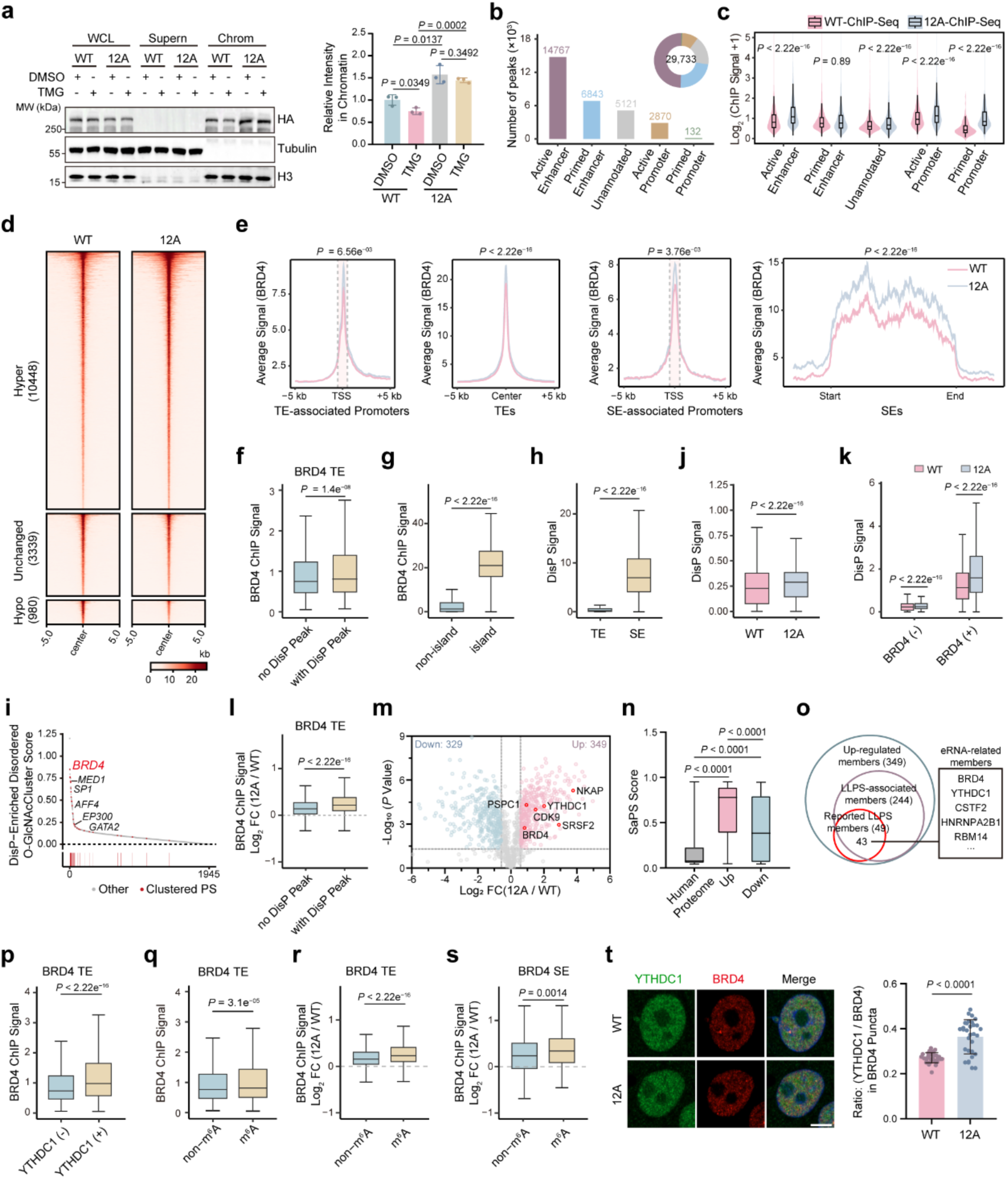
Loss of O-GlcNAc enhances BRD4 chromatin occupancy mostly at active enhancers. **a.** Representative immunoblots and quantitative results showing the amount of chromatin-bound BRD4-WT and BRD4-12A in HeLa cells under indicated treatments. WCL, whole cell lysates. Supern, supernatant. Chrom, chromatin. Quantitative results (right) are the mean ± s.d. from n = 3 biological independent experiments. **b.** Numbers and distributions of BRD4 ChIP-seq peaks across chromatin states marked by histone markers. **c.** Box and violin plots showing the comparisons of BRD4 ChIP-seq signals in BRD4-12A versus BRD4-WT cells across indicated chromatin states. **d.** Heatmaps showing BRD4 ChIP-seq signals at TEs within ± 5 kb in BRD4-12A versus BRD4-WT cells. **e.** Composite plots showing BRD4 ChIP-seq meta-profiles at TEs, SEs and their associated promoters from BRD4-WT and BRD4-12A. Meta-profile centers are located within enhancer regions or transcription start sites, defined as ± 5 kb for TEs and TSSs, and ± 10 kb (including a 3-kb flanking region) for SEs. Plots represent average RPKM per 50 bp bins. TSS, transcription start site; RPKM, Reads Per Kilobase of transcript per Million mapped. Mann-Whitney *U* test with Bonferroni correction was used. **f.** Boxplot showing BRD4 ChIP-seq signals at TEs when matched with or without DisP-seq peaks. **g,** Boxplot showing BRD4 ChIP-seq signals within and outside DisP islands. **h.** Boxplot showing DisP-seq signals at BRD4-occupied TE and SE regions. **i.** Plot showing the DisP-Enriched Disordered O-GlcNAcCluster Score of 1945 ENCODE transcription factors. Experimentally validated phase separating proteins with O-GlcNAc clusters were highlighted in red dots. **j.** Boxplot showing DisP-seq signals of BRD4-WT and BRD4-12A cells. **k.** Boxplot showing DisP-seq signals within or outside BRD4-occupied regions defined by BRD4 ChIP-seq signal in BRD4-WT and BRD4-12A cells. BRD4 (-), non-BRD4-occupied regions. BRD4 (+), BRD4-occupied regions. **l,** Boxplot showing comparison of fold changes of BRD4 ChIP-seq signal in BRD4-12A versus BRD4-WT cells between sites with or without DisP-seq peaks. **m.** Volcano plot showing identified chromatin-bound proteins in BRD4-12A versus BRD4-WT cells. Proteins are color-coded as follows: light pink (log_2_ FC > 0.58), light blue (log_2_ FC < -0.58), and gray (no significant changes). Representative proteins were annotated in red. **n,** Box plot showing SaPS scores of the upregulated and the downregulated proteins in **m** compared to the human proteome. **o.** Venn diagram showing the overlaps among the upregulated proteome (349), LLPS-associated members (244), and the reported LLPS members (49). Representative proteins relating to eRNA and enhancers are listed. **p,** Boxplot showing BRD4 ChIP-seq signal at or outside YTHDC1-occupied regions defined by YTHDC1 ChIP-seq peaks. YTHDC1 (-), non-YTHDC1-occupied regions; YTHDC1 (+), YTHDC1-occupied regions. **q.** Box plot showing BRD4 ChIP-seq signal at m^6^A-modified regions and non-m^6^A modified regions in BRD4-occupied TEs. **r, s.** Box plots showing comparison of fold changes of BRD4 ChIP-seq signals in BRD4-12A versus BRD4-WT cells between m^6^A-modified sites and non-m^6^A-modified sites at TEs (**r**) and SEs (**s**). **t.** Representative immunofluorescence images and quantified results showing the localization and fluorescence density ratio of YTHDC1 and BRD4 in BRD4 puncta at WT and 12A cells. n = 31 and 32 cells in WT and 12A cells, respectively, pooled from three independent replicates. Scale bar, 5 μm. FC, fold change. Data in **a, t** are the mean ± s.d.. Two-tailed t-tests were used for statistical analyses in **a, t**, two-sided Wilcoxon rank-sum tests were used for statistical analyses in **c, e–h, j–l, n, p–s**.

To determine whether this effect stems from O-GlcNAc-mediated modulation of BRD4 phase separation, we performed DisP-seq^47^—a method for mapping IDR-dependent chromatin interactions—in both BRD4-rescue HeLa cell lines (**Extended Data** Fig. 6d). Notably, DisP-seq signal was strongly correlated with ChIP-seq signal at BRD4-bound TEs, and DisP-marked TEs possessed higher BRD4 occupancy than non-DisP-marked TEs (**Fig. 5f, Extended Data** Fig. 6e). These data emphasize the contribution of phase separated BRD4 for chromatin binding. DisP islands, referring to a set of large DisP-seq clusters, were found to concentrate disordered proteins and be associated most with active enhancers. Accordingly, we identified 1,065 DisP islands (**Extended Data** Fig. 6f) and observed that BRD4 ChIP-seq signal was significantly higher within DisP islands than outside (**Fig. 5g**), suggesting that BRD4 is concentrated in DisP islands for regulatory potential to active enhancers. Given that BRD4 is more enriched in SEs relative to TEs, we also examined DisP-seq signal at both BRD4-bound TEs and SEs, and observed a dramatically stronger signal at SEs than TEs (**Fig. 5h**), consistent with SEs being particularly sensitive to condensate perturbations due to their reliance on coactivator phase separation.

We further developed a composite DisP score incorporating peak overlap and O-GlcNAc cluster density for 1,945 ENCODE transcription factors. BRD4 ranked highest among LLPS-prone factors (**Fig. 5i**), underscoring the role of IDR O-GlcNAc clusters in modulating LLPS-dependent chromatin association. Consistently, BRD4-12A-expressing cells exhibited stronger genome-wide DisP-seq signals, with more upregulated peaks compared to BRD4-WT (**Fig. 5j, Extended Data** Fig. 6g). Moreover, matching DisP-seq and ChIP-seq signals revealed that BRD4-12A exhibited stronger DisP-seq signals at BRD4-occupied loci than BRD4-WT (**Fig. 5k**), highlighting that BRD4 IDR-mediated phase separation is linked to chromatin occupancy, with BRD4-12A possessing a better PS capability. At BRD4-bound TEs, DisP-marked TEs exhibited larger fold-change between BRD4-12A and BRD4-WT (**Fig. 5l**), confirming that these loci are sensitive to O-GlcNAc-mediated phase separation perturbation. Among TEs, hyper-binding sites in BRD4-12A versus BRD4-WT showed the most enriched DisP peaks and highest DisP-seq signals (**Extended Data** Fig. 6h, i). Collectively, these findings demonstrated that O-GlcNAc clusters at BRD4 IDR affects BRD4 binding across the genome, in particularly at active enhancers, by tuning BRD4 PS capability.

### BRD4’s chromatin recruitment through YTHDC1 is regulated by O-GlcNAcylation

Given that loss of O-GlcNAcylation enhances BRD4 chromatin occupancy, we next sought to identify the factors responsible for BRD4 recruitment to chromatin and to clarify how this process relates to O-GlcNAc–modulated BRD4 phase separation. We first performed chromatin-associated proteomics in BRD4-WT and BRD4-12A rescue HeLa cells using a crosslinking-based mass spectrometry approach^48^ and focused on proteins showing differential chromatin-binding intensities (**Extended Data** Fig. 7a). Volcano plot analysis revealed 349 upregulated and 329 downregulated nuclear proteins exhibiting more than 1.5-fold change in BRD4-12A cells relative to BRD4-WT cells (**Fig. 5m**), with BRD4 itself showing consistently elevated chromatin binding in BRD4-12A cells. GO analyses of upregulated proteins revealed enrichment for transcription-related pathways and included many transcription factors, epigenetic modulators, and splicing factors (**Extended Data** Fig. 7b-d).

We next asked whether the observed chromatin-binding differences were associated with BRD4’s phase separation propensity. Although the proportion of predicted phase separating proteins was similar in both upregulated and downregulated groups (**Extended Data** Fig. 7e), the upregulated proteins exhibited substantially higher average SaPS scores, indicating stronger LLPS potential (**Fig. 5n**). We speculate that phase separating proteins may facilitate the chromatin recruitment of BRD4. 43 of the upregulated protein are experimentally validated LLPS proteins (**Fig. 5o**). Among them, we further focused on factors known to associate with enhancers, enhancer RNAs (eRNAs) and eRNA-driven condensation. In particular, we noted YTHDC1, which is known to bind methylated-eRNAs and promote condensate formation^49^. We examined the binding of YTHDC1 at BRD4-occupied TEs and SEs, and found a strong correlation between them (**Extended Data** Fig. 7f, g). BRD4 also showed higher binding intensity at genomic loci co-occupied by YTHDC1 (**Fig. 5p**). We thus hypothesized that YTHDC1 may bind to m^6^A-modified eRNAs and facilitate BRD4 recruitment to chromatin. Consistently, we found that YTHDC1 preferentially binds to m^6^A-eRNA sites over non-m^6^A-eRNA sites (**Extended Data** Fig. 7h), and BRD4 showed corresponding enrichment at m^6^A-eRNA sites (**Fig. 5q**), supporting the notion that YTHDC1 promotes BRD4 recruitment. This preference was also observed at SEs, which harbor more m^6^A peaks than TEs (**Extended Data** Fig. 7i-k).

We next assessed whether YTHDC1-mediated BRD4 recruitment is sensitive to O-GlcNAc-mediated modulation of BRD4 phase separation. Genomic regions with increased BRD4 binding in BRD4-12A versus BRD4-WT cells were enriched for YTHDC1 signal (**Extended Data** Fig. 7l), and showed larger differences at m^6^A-eRNA sites than at non-m^6^A-eRNA sites (**Fig. 5r and Extended Data** Fig. 7m). Similarly, at SEs, BRD4 binding at m^6^A-eRNA sites was more sensitive to O-GlcNAcylation, with BRD4-12A exhibiting stronger occupancy than BRD4-WT (**Fig. 5s**). These results support a model in which YTHDC1 facilitates BRD4 recruitment to m^6^A-eRNA sites through a co-condensation mechanism that is sensitive to BRD4 O-GlcNAcylation. Further supporting this model, we observed increased chromatin binding of YTHDC1 in BRD4-12A cells (**Extended Data** Fig. 7n) and confirmed that YTHDC1 colocalizes with BRD4 in nuclear puncta, with greater enrichment in BRD4-12A rescue cells (**Fig. 5t**), suggesting a functional association with BRD4’s enhanced phase separation capacity. *In vitro* droplet assays further demonstrated that YTHDC1 partitions into BRD4 condensates, which are attenuated by BRD4 O-GlcNAcylation (**Extended Data** Fig. 7o). Together, these results support a model in which m^6^A-eRNAs recruit YTHDC1 to active enhancers, further promoting BRD4 condensate assembly. This cooperative interaction enhances BRD4 chromatin binding in BRD4-12A cells and illustrates how transcriptional condensates are sensitive to modulation by O-GlcNAcylation.

### O-GlcNAcylation of BRD4 modulates transcription and cellular fitness

In addition to higher SaPS scores, we also noticed that the upregulated chromatin-binding proteins in BRD4-12A cells versus BRD4-WT cells contain more transcription cofactors than downregulated proteins (**Extended Data** Fig. 8a), including 18 proteins that have been experimentally validated to undergo LLPS (**Extended Data** Fig. 8b). And 9 of 18 LLPS proteins are known to interact with Pol II. Among these, PSPC1 is a paraspeckle protein known to promote Pol II CTD phase separation and phosphorylation via phase separation, thereby facilitating active gene expression^48^. Immunofluorescence imaging showed that PSPC1 colocalized with BRD4 nuclear condensates, with stronger PSPC1 enrichment observed in BRD4-12A cells (**Extended Data** Fig. 8c). Similar to YTHDC1, O-GlcNAcylation of BRD4 reduced its co-condensation with PSPC1 *in vitro* (**Extended Data** Fig. 8d). Given those, we hypothesized that BRD4-12A cells—with greater BRD4 phase separation capability—would exhibit increased transcriptional activity. Supporting this, EU labeling assays revealed a significant elevation in nascent RNA synthesis in BRD4-12A cells compared to BRD4-WT cells (**Fig. 6a**). Consistently, the levels of three histone modifications associated with active transcription—H3K27ac, H3K4me1, and H3K4me3— were elevated in BRD4-12A cells and dynamically responded to BRD4 O-GlcNAcylation status (**Extended Data** Fig. 8e–h). These results indicate that loss of O-GlcNAcylation on BRD4 enhances enhancer activity and transcription. To examine genome-wide transcriptional activity, we analyzed chromatin-associated RNA (caRNA), which reflects pre-mRNA abundance and closely correlates with transcriptional output^50^. Notably, there was a strong positive correlation between caRNA changes and BRD4 chromatin occupancy, with most genes showing both increased BRD4 binding and higher transcription levels in BRD4-12A cells (**Fig. 6b**). Similar correlations were observed between BRD4 binding and nuclear or cytoplasmic mRNA levels (**Extended Data** Fig. 8i), suggesting that BRD4 chromatin occupancy influences nearby gene expression, likely through transcriptional regulation. Gene set enrichment analysis (GSEA) revealed that genes with over 1.5-fold changes in BRD4 binding in BRD4-12A versus BRD4-WT cells exhibited global upregulation at the total RNA level (**Extended Data** Fig. 8j). SEs contain genes with the highest upregulation in BRD4-12A cells, followed by TEs (**Fig. 6c** and **Extended Data** Fig. 8k). GSEA also confirmed that SE-associated genes were significantly upregulated in BRD4-12A cells (**Fig. 6d**), indicating that loss of O-GlcNAcylation leads to increased transcriptional activity, particularly at SEs.

**Fig. 6.**
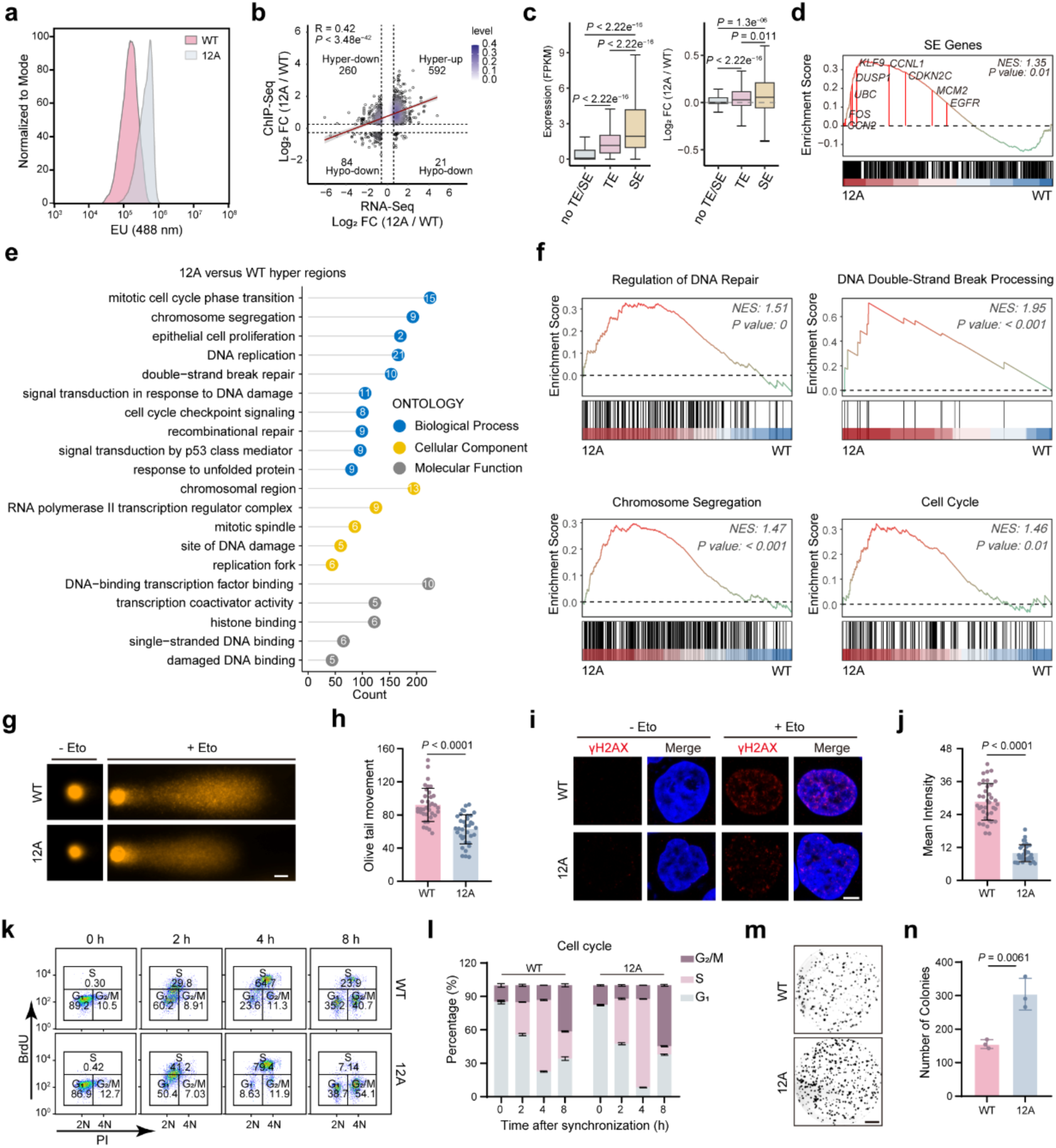
Loss of O-GlcNAc on BRD4 promotes transcription and affects pathways relating to DNA damage, cell cycle and proliferation. **a.** Flow cytometric analysis of EU labelling showing nascent RNA levels in BRD4-12A and BRD4-WT cells. EU, 5-Ethynyluridine. **b.** The correlation between fold changes of BRD4-12A versus BRD4-WT in chromatin RNA-seq and in ChIP-seq. Black circles and the purple gradient represent single genes and gene densities, respectively. Two-sided Pearson’s correlation coefficients (R) with *P* values were used. **c.** Box plot showing gene expression levels (left), as well as gene expression fold changes of BRD4-12A versus BRD4-WT at TEs, SEs and other no TE/SE sites (right). FC, fold change. Two-sided Wilcoxon rank-sum tests were used for statistical analyses. **d.** Gene set enrichment analysis (GSEA) showing SE-related gene expression differences between BRD4-WT and BRD4-12A cells. Representative genes were annotated. **e.** Gene Ontology (GO) analysis of genes with increased ChIP-seq intensities in BRD4-12A versus BRD4-WT cells. GO terms associated with molecular function, cellular component, and biological processes are highlighted in gray, yellow, and blue circles with -log_10_ *P* inside, respectively. **f.** GSEA plots showing relevant pathways with differential gene expression in BRD4-12A cells versus BRD4-WT cells. NES, normalized enrichment score. **g, h.** Representative images (**g**) and quantification results (**h**) of alkaline comet single cell gel electrophoresis assays performed on BRD4-WT and BRD4-12A cells. n = 36 and 35 cells in WT and 12A groups, respectively, pooled from three biological independent replicates. **i, j.** Representative immunofluorescence images (**i**) and quantification (**j**) of γH2AX intensities in BRD4-WT and BRD4-12A cells. n = 37 and 35 cells in WT and 12A groups, respectively, pooled from three independent replicates. **k, l.** Flow cytometric cell cycle analyses (**k**) and quantification results (**l**) of BRD4-WT and BRD4-12A cells at indicated time points post release from cell synchronization by double thymidine**. m, n.** Colony formation analysis (**m**) and quantification results (**n**) of BRD4-WT and BRD4-12A cells. n = 3 biologically independent experiments. Scale bar, 20 μm (**g**), 5 μm (**i**) and 5 mm (**m**). Data in **h, j, l, n** are the mean ± s.d.. Two-tailed t-tests were used for statistical analyses in **h, j, n**.

To better understand the functional implications, we conducted GO analysis of regions with elevated BRD4 binding in BRD4-12A cells and found enrichment in pathways related to the cell cycle, DNA damage repair, and proliferation (**Fig. 6e**). GSEA further confirmed that genes involved in these pathways were significantly upregulated in BRD4-12A cells (**Fig. 6f**), in line with chromatin RNA-seq results (**Extended Data** Fig. 9a). BRD4 is well known to participate in regulating the cell cycle, proliferation, and DNA repair, particularly in cancer contexts^51^. Representative genes from these enriched pathways—including *H2AX*, *TGFB1*, and *CCN2*—showed higher signal intensities in both ChIP-seq and RNA-seq in BRD4-12A cells (**Extended Data** Fig. 9b). ChIP-qPCR and RT-qPCR validated increased BRD4 binding and mRNA expression for eight selected genes in BRD4-12A compared to BRD4-WT cells (**Extended Data** Fig. 9c, d). These data support the conclusion that O-GlcNAcylation on BRD4 suppresses gene expression linked to critical cellular processes by limiting BRD4 chromatin occupancy.

We next assessed whether these transcriptional changes led to functional phenotypes. To evaluate DNA damage response, we treated cells with etoposide (Eto)^52^, a topoisomerase II inhibitor, and measured DNA damage using comet assays and γH2AX immunostaining. After Eto treatment, BRD4-WT cells displayed longer comet tails (**Fig. 6g, h**) and higher γH2AX foci intensity (**Fig. 6i, j**) than BRD4-12A cells, suggesting that O-GlcNAcylation sensitizes cells to DNA damage. We also examined cell cycle progression and found that BRD4-12A cells exhibited significantly faster G1-to-S phase transition following release from synchronization, consistent with unsynchronized cell cycle profiles (**Fig. 6k, l, Extended Data** Fig. 9e, f). Colony formation assays further confirmed that BRD4-12A cells proliferated more rapidly than BRD4-WT cells (**Fig. 6m, n**). Collectively, these results establish a functional link between BRD4 O-GlcNAcylation and transcriptional repression of genes involved in cell cycle progression, proliferation, and DNA repair. Loss of O-GlcNAcylation enhances BRD4 chromatin binding and transcriptional activity, ultimately promoting cell proliferation and stress resistance. These findings highlight O-GlcNAcylation as a critical regulatory mechanism that tunes BRD4-mediated transcription and cell fitness.

## Discussion

Considerable effort has gone into delineating the molecular principles that govern the biophysical properties of biomolecular condensates, with PTMs receiving growing attention because of their dynamic and reversible nature, as well as diverse chemical attributes^15–17^. As one of the most abundant PTMs, O-GlcNAc has been implicated in modulating protein phase separation in several cases^32–34^, yet systematic studies remain scarce. In this study, by interrogating the human O-GlcNAcome, we uncovered a strong correlation between O-GlcNAc sites and IDRs, and linked this relationship to PS propensity (**Fig. 1**). Although the global O-GlcNAcome lacks a defined consensus motif, IDRs show unexpected enrichment of clustered Pro, Ser and Thr residues that host O-GlcNAc clusters. The presence of O-GlcNAc clusters were strongly associated with protein PS propensity. This comprehensive analysis provides the first proteome-wide evidence that clustered O-GlcNAc marks on IDRs of phase-separating proteins, adding a new regulatory layer to protein phase separation.

Although O-GlcNAcylation is often sub-stoichiometric, the formation of O-GlcNAc clusters likely compensates for low occupancy at individual sites, resulting in a cumulative effect. In BRD4, for example, none of the twelve glycosylation sites is individually essential (**Fig. 2**), yet their collective modification alters condensate biophysics by increasing local hydrophilicity and steric hindrance, thereby reducing phase separation capacity and stability (**Fig. 3, 4**). We therefore anticipate similar modulatory roles in LLPS across other proteins bearing O-GlcNAc clusters, especially transcription-related proteins. In addition to O-GlcNAc clusters, site-specific O-GlcNAcylation may regulate LLPS and aggregation by altering local structure or disrupting interaction interfaces^32^. Although such site-specific effects remain mostly confined to case studies, how O-GlcNAc clusters coordinate with individual glycosites to modulate condensate behavior warrants further exploration. Notably, similar cluster-based modulation has been observed for other PTMs, such as multisite phosphorylation and PARylation in Ki-67^21^ and CycT1^24^, further supporting the concept that clustered PTMs in IDRs represent a general mechanism for LLPS regulation by accumulating subtle chemical changes across extended regions. Our proteome-wide analysis of O-GlcNAc patterns within IDRs offers a framework for investigating other PTMs and refining the “molecular grammar” that governs condensate behavior.

O-GlcNAcylation has long been implicated in the transcriptional machinery^53,54^, but its functional mechanisms remain incompletely understood. Here, we show that O-GlcNAc clusters are particularly enriched in transcriptional regulators—including chromatin remodelers and coactivators—that are likely to undergo IDR-driven phase separation for transcriptional modulation. Using BRD4 as a model, we demonstrate that O-GlcNAcylation attenuates its chromatin binding by reducing its PS propensity. ChIP-seq and chromatin proteomics analyses revealed that preventing BRD4-IDR O-GlcNAcylation significantly enhances BRD4 occupancy at active enhancers, with SEs being especially affected (**Fig. 5**). DisP-seq analyses further confirmed that the observed changes in BRD4 chromatin binding and transcriptional activity are driven by condensate perturbation. Chromatin engagement of transcriptional factors and chromatin remodelers is critical to downstream transcriptional activities, which is typically attributed to structured DNA-binding or histone-binding domains and sequence-specific recognition of motifs. However, our findings reveal a distinct mechanism whereby O-GlcNAc clusters within IDRs—regions not directly responsible for DNA recognition—modulate chromatin binding through condensate dynamics. This O-GlcNAc-dependent regulation extends beyond BRD4, where we observed coordinated changes in chromatin association for other LLPS-prone regulators, highlighting a broader transcriptional rewiring driven by condensate modulation. Among these, we identified YTHDC1 as a co-regulator that promotes BRD4 recruitment to active enhancers in an m^6^A-eRNA–dependent manner. While YTHDC1 exemplifies one such pathway, additional factors are likely to intersect with O-GlcNAc-modified BRD4 condensates to influence final transcriptional outputs.

The unique phase separation properties at SEs render them highly sensitive to condensate perturbation^55^. Consistently, SE-associated genes were the most responsive to O-GlcNAcylation-induced changes in BRD4 condensation, displaying reduced transcription in conjunction with diminished BRD4 binding. As a central player at SEs, BRD4 perturbation led to widespread transcriptional changes across hundreds of SEs, affecting key cellular programs such as DNA damage repair, cell cycle progression, and proliferation (**Fig. 6**). These findings emphasize the importance of uncovering new regulatory mechanisms underpinning key factors within transcriptional condensates. Despite the inherent challenges posed by the dynamic and fragile nature of transcriptional condensates and the sometimes-low PTM stoichiometry, future studies can explore how these dynamically modified substrates cooperatively orchestrate transcriptional processes across different biological contexts from a systemic perspective^54,56^.

In summary, O-GlcNAc clusters within IDRs emerges as a widespread cue for regulating phase separation. In BRD4, O-GlcNAc clusters weaken condensate formation, curtail enhancer binding and reprogram gene networks that sustain genome integrity and proliferation (**Fig. 7**). This link between O-GlcNAc modification on BRD4 C-terminal IDR and transcriptional condensates offers new therapeutic angles beyond pharmacological BRD4 inhibition or degradation.

**Fig. 7.**
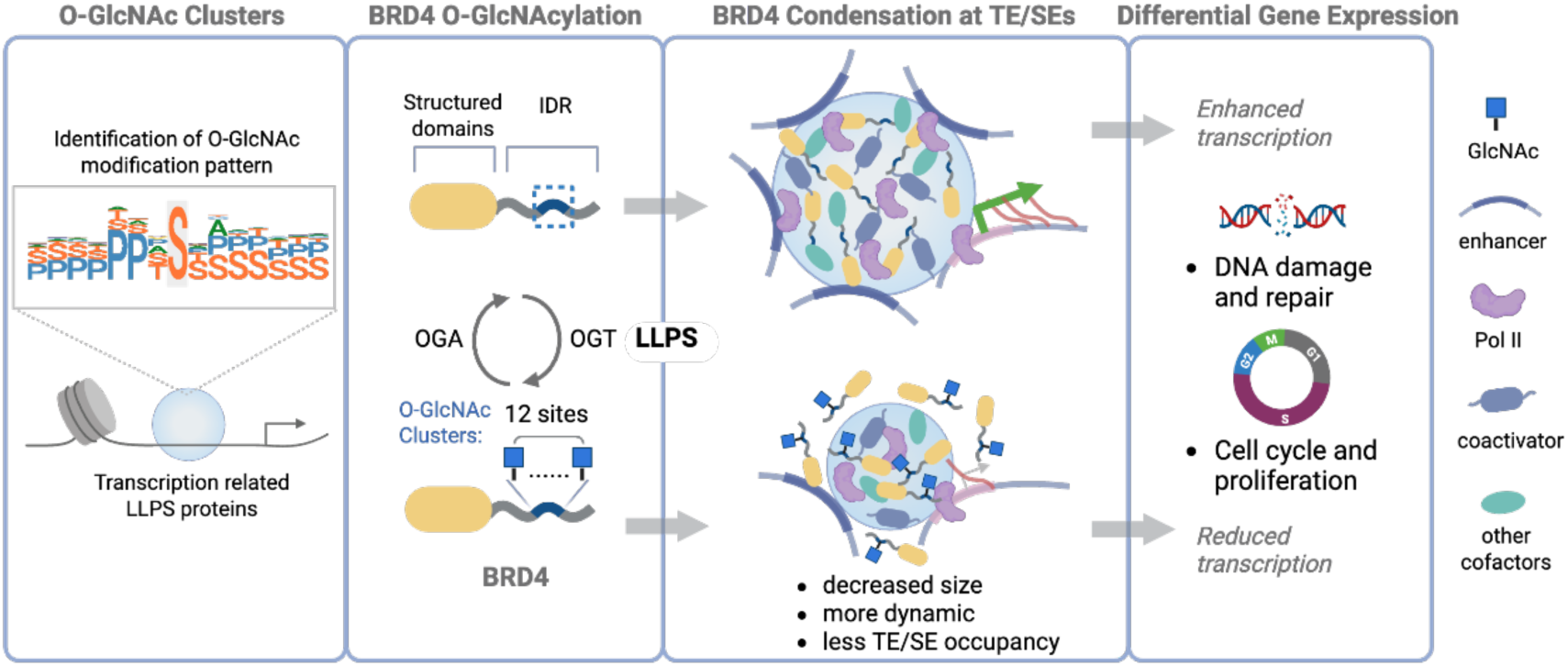
A proposed model of O-GlcNAc clusters on BRD4 regulating its phase separation for gene control. O-GlcNAc clusters are associated with proteins exhibiting LLPS propensity, especially those involved in transcription. Among them, BRD4 was identified with O-GlcNAc clusters at its C-terminal IDR. O-GlcNAc clusters suppress BRD4 phase separation both in vitro and in cells. Reduced BRD4 condensates by O-GlcNAcylation lead to less occupancy at SEs and thereby reduced transcriptional activities on pathways associated with DNA damage and repair, as well as cell cycle and proliferation. Representative symbols are shown on the right.

## Acknowledgements

This study is supported by the National Natural Science Foundation of China (92478103 and 22277080 to Y.G. and 32170595 to J.L.), the Major Program of Shenzhen Bay Laboratory (S241101001), the Startup Fund from Shenzhen Bay Laboratory, and the Beijing Nova Program to J.L. (Z211100002121011 and 20230484442). We thank the Multi-Omics Mass Spectrometry Core of the Biomedical Research Core Facilities (BRCF) and the Bioimaging Core at Shenzhen Bay Laboratory, and the Core Facilities of Life Sciences at Peking University for technical support.

## Author contributions

Y.G. and J.L. conceived the idea and designed the original studies. G.S. collected most data except sequencing and bioinformatics, with assistance from C.L., N.W., and Y.G.. T.W. conducted the sequencing experiments and Q.D. performed the bioinformatics analyses. Y.G. and J.L. wrote the paper with input from all authors.

## Methods

### Cell culture and generation of stable cell lines

HeLa cells and HEK293T were cultured at 37 °C with 5% CO_2_ in Dulbecco’s modified Eagle’s medium (DMEM) (Gibco) supplemented with 1% penicillin/streptomycin (P/S) (Gibco) and 10% FBS (Gibco).

To generate a stable *BRD4* knockdown HeLa cell line, we inserted shRNA targeting *BRD4* (5’-CAGTGACAGTTCGACTGATGA-3’) into the pLKO.1-shRNA plasmid. Lentiviruses were produced by co-transfection of pLKO.1-sh*BRD4*, psPAX2 and pMD2.G into HEK293T cells and collected 48 hours after transfection. HeLa cells were infected for 24 h and then selected with 2 μg/ml puromycin (InvivoGen, ant-pr-1) for over two days.

To rescue sh*BRD4* HeLa cells with EGFP-BRD4-WT or EGFP-BRD4-12A, the pPB-CAG-BRD4-blasticidin plasmid was constructed by cloning BRD4 or B12A cDNAs into a PiggyBac Transposon System (SBI). The plasmid was transfected into sh*BRD4* HeLa cells. After 24 h, HeLa cells were subjected to selection with 2 μg/ml blasticidin (InvivoGen, ant-bl-1) for over two days. EGFP-positive cells were sorted by CytoFLEX SRT (BACKMAN COULTER). BRD4 knockdown and rescue cells were confirmed using immunoblot and RT-qPCR.

### Protein expression and purification

N-terminal EGFP-fused BRD4 variants and N-terminal mCherry-fused YTHDC1 and PSPC1 were subcloned into the pET-30a(+) vector containing 6× His tags. His-tagged OGT and its inactive mutant OGT (K852A) were subcloned into the pET24b(+) vector. These plasmids were transformed into *E. coli.* DE3 (BL21).

After induction with 1 mM Isopropyl-β-D-thiogalactoside, bacteria lysates in buffer (20 mM Tris pH 7.5, 500 mM NaCl and 1 mM PMSF) were sonicated, and the His-tagged proteins were purified using the 5 mL HisTrap™ HP (Cytiva, 17524802). The eluted proteins were concentrated using Amicon Ultra-15 spin columns (Merck-Millipore, 30K) into corresponding storage buffer: BRD4-IDR storage buffer (50 mM Tris pH 7.5, 500 mM NaCl, 10% glycerol, 1 mM DTT); OGT storage buffer (25 mM Tris pH 7.5, 50 mM NaCl, 0.5 mM EDTA, 10% glycerol, 2 mM DTT); PSPC1 storage buffer (20 mM Tris pH 8.0, 150 mM NaCl, 10% glycerol); YTHDC1 storage buffer (20 mM HEPES pH 7.4, 300 mM KCl, 6 mM MgCl_2_, 0.02% NP-40, 10% glycerol).

### *In vitro* glycosylation assay

For each reaction, 10 µM purified substrate protein, 1 mM UDP-GlcNAc (Millipore, U4375), and 1 µM purified OGT were mixed as indicated in the OGT storage buffer. The reaction was conducted at 30 °C for 4 h with rotation. Samples were concentrated into the BRD4-IDR storage buffer at 4 °C using Amicon Ultra-0.5 spin columns (Millipore, 3K) and stored at -80 °C after determining protein concentrations.

### Immunoprecipitation and immunoblot assays

According to the manufacturer’s instructions of anti-HA magnetic beads (Selleck, B26202), cells with indicated treatments were lysed with RIPA lysis buffer containing 1× Protease Inhibitor Cocktail (EDTA-free) (MCE, HY-K0010) and 20 μM Thiamet-G (TMG, Selleck, S7213). Cell lysates were incubated with prewashed anti-HA magnetic beads with gentle rotation at 4 °C overnight. After washed three times with 1× PBST buffer (137 mM NaCl, 2.7 mM KCl, 10 mM Na_2_HPO_4_, 1.8 mM KH_2_PO_4_, 0.5 % Tween-20), proteins were eluted by 2× SDS loading buffer at 95 °C for 10 min.

For immunoprecipitation of endogenous BRD4, cells were lysed in lysis buffer (50 mM HEPES pH 7.5, 150 mM KCl, 2 mM EDTA, 0.5% NP40) containing freshly added 0.5 mM DTT, complete EDTA-free protease inhibitor cocktail and 20 μM TMG. Cell lysates were incubated with Pierce™ Protein A Magnetic Beads (Thermo Scientific, 88845) or Pierce™ Protein G Magnetic Beads (Thermo Scientific, 88847), which were pre-incubated with anti-BRD4 antibody at 4 °C overnight. After washed three times using the wash buffer (50 mM HEPES–KOH pH 7.5, 300 mM KCl, 0.05 % NP40, 0.5 mM DTT, protease inhibitor cocktail and 20 μM TMG), proteins were eluted with SDS loading buffer for the following immunoblot experiment.

Unless otherwise noted, proteins were analyzed using the precast SDS-PAGE gel and transferred to a nitrocellulose membrane (Millipore, HATF00010). The membrane was blocked with 5% BSA/PBST. Proteins were detected by the indicated primary and HRP or IRDye secondary antibodies. Images were acquired by Azure Imager C600 (Azure Biosystems) and analyzed using Fiji ImageJ.

Primary and secondary antibodies used for immunoprecipitation, flow cytometry, ChIP-seq and ChIP-qPCR, immunoblot, and Immunofluorescence include: mouse anti-BRD4 (Cell Signaling, 63759; 1:1,000 for immunoblot, and 1:100 for immunoprecipitation, and 1:1600 for immunofluorescence), rabbit anti-BRD4 (Cell Signaling, 13440; 1:50 for ChIP-seq and ChIP-qPCR), rabbit anti-BRD4 (Abcam, ab243862; 1:200 for immunofluorescence), mouse anti-O-GlcNAc (RL2) (Abcam, ab2739; 1:1,000 for immunoblot), rabbit anti-GAPDH (HUABIO, ET1601; 1:2,000 for immunoblot), mouse anti-His_6_ (HUABIO, HA601079; 1:1,000 for immunoblot), rabbit anti-HA (Cell Signaling, 3724; 1:1000 for immunoblot; 1:50 for ChIP-seq and ChIP-qPCR), mouse anti-Myc (Cell Signaling, 2276; 1:1000 for immunoblot), mouse anti-GFP (Abclonal, AE012; 1:5000 for immunoblot), rabbit anti-OGT (Cell Signaling, 24083; 1:1000 for immunoblot), rabbit anti-YTHDC1 (Abcam, ab122340; 1:500 for immunoblot, 2 μg/ml for immunofluorescence), mouse anti-PSPC1 (Santa Cruz, sc-374181; 1:100 for immunoblot, 1:200 for immunofluorescence), mouse anti-BrdU (Cell Signaling, 5292; 1:200 for flow cytometry), rabbit anti-β-Tubulin (Servicebio, GB11017; 1:1000 for immunoblot), rabbit anti-Histone H3 (Cell Signaling, 4499; 1:2000 for immunoblot), rabbit anti-Phospho-Histone H2A.X (Ser139) (Cell Signaling, 9718; 1:400 for immunofluorescence), rabbit anti-Histone H3 (acetyl K27) (Abcam, ab4729; 1 µg/ml for immunoblot), rabbit anti-Histone H3 (mono-methyl K4) (Cell Signaling, 5326; 1:1000 for immunoblot), rabbit anti-Histone H3 (tri-methyl K4) (Cell Signaling, 9751; 1:1000 for immunoblot), HRP-conjugated antibody to biotin (Cell Signaling, 7075; 1:1000 for immunoblot), HRP-conjugated secondary antibodies (HUBIO, HA1006 (anti-mouse IgG) or HA1001 (anti-rabbit IgG); 1:10,000 for immunoblot), anti-Mouse Alexa Fluor™ 488 secondary antibody (Invitrogen, A11001; 1:1000), anti-Mouse Alexa Fluor™ 568 secondary antibody (Invitrogen, A11004; 1:1000), anti-Rabbit Alexa Fluor™ 488 secondary antibody (Invitrogen, A11008; 1:1000), anti-Rabbit Alexa Fluor™ 568 secondary antibody (Invitrogen, A11011; 1:1000), anti-Mouse Alexa Fluor™ 647 (Invitrogen, A21235; 1:1000).

### Chemoenzymatic labeling and enrichment of O-GlcNAcylated proteins

The proteins enriched by anti-HA magnetic beads were labeled with GalNAz as previously reported^57^. Briefly, immunoprecipitated beads were incubated with GalT1 (Y289L), UDP-GalNAz, MnCl_2_ in the GalT1 labeling buffer (50 mM NaCl, 20 mM HEPES, 2 % NP-40, pH 7.9) at 4 °C for at least 20 h with gentle rotation. Labeled proteins were then clicked with 200 µM Biotin-Alkyne (Confluore, BBBE-1) in the presence of 0.4 mM CuSO_4_, 2.5 mM fresh sodium ascorbate and 0.8 mM BTTAA at 30 °C for 1 h. The proteins were eluted with SDS loading buffer and analyzed by immunoblot.

### Identification of BRD4 O-GlcNAc sites

Following *in vitro* O-GlcNAcylation, 20 μg of the product were denatured with 6 M urea in 50 mM Tris pH 8.0, then sequentially reduced and alkylated with 10 mM tris-(2-carboxyethyl)-phosphine (TCEP) (Millipore, C4706) for 30 min and 10 mM 2-Iodoacetamide (IAA) (Millipore, I6125) for 45 min at room temperature. Excess IAA was quenched with 10 mM TCEP. The sample was diluted to 0.8 mM urea with 50 mM Tris pH 7.4, 1 mM CaCl_2_ and digested with trypsin (Promega, V5111) or chymotrypsin (Promega, V1061) overnight. The resulting peptides were desalted with ZipTip (Millipore, ZTC18S096) and concentrated to dryness by Eppendorf Vacufuge for storage at -80 °C until analysis.

Glycosites were identified using a Thermo Scientific Orbitrap Fusion Lumos Tribrid mass spectrometer with an EASY-nLC 1000 system and a nano-electrospray ion source. The instrument parameters were set according to previous reports unless otherwise noted. The raw data was processed using PMI Byonic™ and pGlyco3 by searching against the EGFP-fused BRD4-IDR protein sequence with the following settings: trypsin or chymotrypsin as enzyme, 2 missed cleavages allowed; 10 ppm precursor tolerance and 20 ppm fragment tolerance on fragment ions; cysteine carbaminomethylation as a fixed modification; oxidation on M and deamidated N/Q as variable modifications. For Byonic, HexNAc on S/T was set as a variable modification; peptides allow a maximal charge of 4. For pGlyco3, ‘HCD+ETHCD’ was selected as the Fragmentation method; Multi-Site-O-Glycan database was used with glycosylation sites at S/T. Other parameters followed the default settings. All spectra of peptides were displayed using the pLabel (version 2.4).

### Identification of the chromatin-bound proteome

Chromatin binding of BRD4, PSPC1 and YTHDC1 was detected following a previously reported procedure^58^. In brief, BRD4-WT or BRD4-12A HeLa cells were seeded in a 10 cm dish with indicated treatments. After 48 h, cells were harvested and lysed with a chromatin extraction buffer (20 mM Tris-HCl, 100 mM NaCl, 5 mM MgCl_2_, 10% glycerol, 0.2% IGEPAL CA-630, 2 mM NaF, 2 mM Na_3_VO_4_, Protease Inhibitor Cocktail) for 1 h at 4 °C, with 5 s vortex every 10 min. 10 μL of the cell lysate was used as the whole cell lysate (WCL) fraction. The remaining cell lysate was centrifuged using 16,000 g for 5 min at 4 °C. The supernatant was collected, and the insoluble fraction was washed three times with the chromatin extraction buffer. All fractions were digested with Benzonase (Sigma, E1014) and then supplemented with 0.1% SDS. After further sonication, they were denatured in SDS loading buffer and subjected to immunoblot.

For identification of the chromatin-bound proteome^48^, cells (∼1 × 10⁷) were lysed in 200 μL NP-40 buffer (10 mM Tris-HCl, pH 7.5, 0.15% NP-40, 150 mM NaCl) on ice for 5 min. The lysate was then resuspended with a 500 μL of 24% sucrose cushion and centrifuged at 15,000 g for 10 min at 4°C. The pelleted nuclei were crosslinked with 1% formaldehyde for 10 min, quenched by 0.125 M glycine, and resuspended in twice the pellet volume of nuclei lysis buffer (50 mM Tris-HCl, 10 mM EDTA, 1% SDS). DNA-protein complexes were precipitated with ethanol at -20°C for 1 h, followed by two washes with 75% ethanol. The pellet was solubilized in Tris-HCl buffer (pH 7.4) supplemented with 8 M urea and 2% SDS at 37°C for 30 min, treated with an equal volume of 5 M NaCl for 30 min, and reprecipitated by adding 0.1 volume of 3 M sodium acetate and three volumes of ethanol. The mixture was centrifuged at 5,000 g for 5 min at 4°C. The pellet was then washed twice with 75% ethanol and resuspended in 100 μL DNase digestion buffer (20 mM HEPES pH 7.5, 15 mM NaCl, 6 mM MgCl_2_, 1 mM CaCl_2_, 10% glycerol). After DNase I digestion (10 U, 37°C, 1 h) and EDTA quenching, the sample was centrifuged at 13,000 rpm for 20 min at 4°C. The supernatant was subjected to SDS-PAGE, followed by in-gel digestion and LC-MS/MS analysis. The raw data were analyzed using Proteome Discoverer software (version 2.5, Thermo Scientific) with the Sequest HT search engine with the following search parameters: trypsin as the digest enzyme with up to 2 missed cleavages; carbamidomethyl[C] as fixed modification, oxidation of M and acetyl or Met-loss on protein N termini as variable modifications; 10 ppm precursor tolerance and 20 ppm fragment tolerance for fragment ions; 1% false discovery rate. Identified proteins with at least two unique peptides, annotated nuclear localization at UniProt database and no missing values were further analyzed by label-free quantification.

### *In vitro* transcription of *Pou5f1* enhancer RNA

According to previous reports^43^, the following primers were used to obtain the DNA sequence of *Pou5f1* enhancer RNA: *Pou5f1*-F: TAATACGACTACTATAGGGGGCCTAGACAGCACTCTCCA; *Pou5f1*-R: TGGATCTCTGTGAGTTCAAG. Based on the DNA sequence, *in vitro* transcription was conducted to synthesize *Pou5f1* enhancer RNA using MEGAscrip T7 transcription kit (Invitrogen, AM1333) according to the manufacturer’s instructions. For visualization of *Pou5f1* enhancer RNA, Cy5-labeled UTP (APExBIO, B8333) was added to the reaction at a final concentration of 1 mM. Transcriptional products were purified through the MEGAclear Transcription Clean-Up Kit (Invitrogen, AM1908) and the purified RNA was quantified using NanoDrop (Thermo Scientific).

### *In vitro* droplet formation and imaging

Indicated purified BRD4 proteins were added into a droplet reaction buffer (50 mM Tris-HCl pH 7.5, 10% glycerol, 1 mM DTT) containing 10% PEG8000 (Coolaber, CP8241) with varying NaCl concentrations. In co-condensation experiments, 10 μM BRD4 was mixed with either 5 μM PSPC1 or 2 μM YTHDC1 in the droplet reaction buffer supplemented with 10% PEG8000 with 125 mM NaCl. For imaging of EGFP-fused BRD4-IDR droplets with *Pou5f1* enhancer RNA, experiments were conducted following previous reports with minor modifications. Briefly, 1 μM substrate protein was added into RNA phase separation buffer (10 μM HEPES pH 7.4,150 mM KCl, 3 mM MgCl_2_ and 0.01% NP-40) containing 10% PEG8000. 10 μL of the reaction sample was immediately loaded into a homemade imaging chamber, consisting of a glass slide and a coverslip attached by two parallel strips of double-sided tape. Images were acquired using ZEISS LSM 980 microscope with a 63× objective and analyzed using Fiji Image J.

### Fluorescence recovery after photobleaching (FRAP) analysis

BRD4 condensates *in vitro* or in cells were photobleached using a laser intensity of 100% at 488 nm and the recovery was recorded at the indicated time points using ZEISS LSM 980 microscope. FRAP data were analyzed using Fiji ImageJ. The raw data were exported, and the pre-bleached fluorescence intensity was normalized to 1, while the post-bleached signal was normalized relative to the pre-bleached level.

### Turbidity assay

Following *in vitro* droplet formation, 30 μL of sample was transferred into a 384-well glass-bottom plate (Cellvis, P384-1.5H-N-1) for measurement on a Spark^®^ Multimode Microplate Reader (Tecan) under 350 nm-wavelength light at room temperature. The optical density (OD) was normalized to that of the BRD4-IDR phase separation solution containing either 50 mM NaCl or 5 μM substrate protein.

### Immunofluorescence and live cell imaging

For imaging BRD4 condensates, the indicated cells were seeded on a 4-Chamber Glass Bottom Dish (JingAnBiological, J40204) for 24 h and treated with 20 μM TMG or OSMI-4 (Selleck, S8910) for additional 48 hours, or with 2.5% 1,6-hexanediol (TCI, H0099) for 10 min. Cells were washed twice with PBS and fixed with 4% paraformaldehyde (Leagene, DF0134) in PBS for 15 minutes at room temperature. After another two washes, cells were blocked with the blocking buffer (5% BSA, 0.3% Triton X-100 in PBS) for 1 h at room temperature. Cells were incubated overnight at 4 °C with indicated primary antibodies followed by three washes, and then incubated with corresponding florescent secondary antibodies for 1 h at room temperature.

For γ-H2AX imaging, cells were treated with 10 μM etoposide (Sigma-Aldrich, E1383) at 37 °C for 1 h before sample preparation, and stained with antibodies and NucBlue™ Fixed Cell ReadyProbes (DAPI) (Invitrogen, R37606) before imaging. Images were acquired on a ZEISS LSM 980 microscope with a 100× objective.

For live cell imaging, sh*BRD4* HeLa cells seeded in the imaging chamber were transiently transfected with the indicated plasmids along with indicated small molecule treatments. 48 h post-transfection, cells were stained with NucBlue™ Live ReadyProbes (Hoechst 33342) (Invitrogen, R37605) for 30 min at 37 °C and then imaged using 100× objective of the Olympus SpinSR microscope equipped with a live cell culture system. All images were analyzed using Fiji ImageJ.

### Comet assay

The Comet Assay Kit (Beyotime, C2041S) was employed to assess DNA damage following the manufacturer’s protocol. In brief, cells treated with Eto were collected and resuspended as a single-cell suspension in 0.7% low-melting-point agarose, then spread onto positively charged adhesive glass slides (Beyotime, FSL051). The slides were then fully covered with lysis buffer (included in the kit) for overnight lysis, then subjected to electrophoresis at 25 V for 25 min in alkaline electrophoresis buffer (200 mM NaOH, 1 mM EDTA, pH 8.0) after a 30-minute equilibration, followed by three times washes with 0.4 M Tris-HCl (pH 7.5) for 5 min each and propidium iodide staining. Images were acquired using a Nikon Ti2E fluorescence microscope, and analyzed with Fiji ImageJ. DNA damage was quantified by Olive tail moment (OTM), calculated as OTM = (Tail mean − Head mean) × (Tail DNA% / 100).

### Flow cytometry analysis of cell cycle

Cells were treated with 10 mM BrdU (Beyotime, ST1056) for 30 min before being collected. For synchronization of cells at G1/S phase, cells were treated with 2 mM thymidine (Beyotime, ST1704) for 16 h and released. After 8 h, cells were treated with 2 mM thymidine again for 16 h and released. Cells were harvested at 0, 2, 4, and 8 h post-release. 10 mM BrdU was added for 30 min before collection. Cells were fixed with 70% ice-cold ethanol at -20 °C overnight followed by treatment with 4 M HCl for 20 min at room temperature. Cells were centrifuged at 4,000 r.p.m. for 10 min and then resuspended in the antibody incubation buffer (5% BSA/PBS) for subsequent incubation at room temperature. Finally, 500 µL of 5 mg/mL PI (MCE, HY-D0815) was added to the samples at 4 °C for 30 min. Fluorescence intensity was detected using Attune NxT Flow Cytometer (Invitrogen). Data were analyzed by FlowJo v10.

### Nascent RNA detection via 5-ethynyl uridine (EU) labeling

EU labeling and nascent RNA detection were performed utilizing Click-iT RNA Alexa Fluor 488 imaging kit (Thermo Scientific, C10329) following the manufacturer’s protocols. Flow cytometry was performed using CytoFlex LX Mosaic88 flow cytometer and analyzed using FlowJo v10.

### Colony formation assay

A total of 750 cells per well were seeded in 6-well plates, with the culture medium replaced every three days. After 10 days, cells were fixed with 4% paraformaldehyde at room temperature for 10 min, then washed twice with water, stained with Crystal Violet Staining Solution (Beyotime, C0121), washed with water three times, and imaged by Azure Imager C600 (Azure Biosystems). Colony number was counted using Fiji ImageJ.

### Protein annotations and analyses of O-GlcNAc site level and density

O-GlcNAc sites were extracted from the human O-GlcNAcome database^37^. Transcriptional cofactors were defined according to the previous report^59^. The O-GlcNAc consensus sequences at different regions were generated as previously described^60^, using -7 to +7 amino-acid sequences surrounding O-GlcNAc sites. Experimentally validated phase-separating proteins (designated PS (Reported)) were compiled from three curated databases: PhaSePro, LLPSDB v2.0, and PhaSepDB, and DrLLPS^38–40,61^. For other human proteins, phase separation propensity was assessed using the SaPS score from PhaSePred database^41^. PS (Predicted) denotes proteins with SaPS ≥ 0.5, while Non-PS (Predicted) denotes those with SaPS < 0.5.

IDRs across the human proteome were annotated according to UniProt. The IDR fraction is defined as the ratio of the IDR length to the full protein length. The O-GlcNAc site level refers to the ratio of O-GlcNAc sites to the total amount of S/T residues within a defined region. The O-GlcNAc site density refers to the density of O-GlcNAc sites per amino acid within a defined region independent of amino acid composition. The IDR-O-GlcNAc site enrichment score quantifies the contribution of clustered O-GlcNAc modifications within IDRs, defined as:

> IDR-O-GlcNAc site enrichment score = (*N*_IDR-O-GlcNAc_ × *L*_IDR_)/*L*_non-IDR_

where *N*_IDR-O-GlcNAc_ is the number of clustered O-GlcNAc sites in IDRs, *L*_IDR_ is the total IDR length and *L*_non-IDR_ is the non-IDR region length of the protein.

### Real-time fluorescence quantification PCR (RT-qPCR)

Total RNA and RNA associated with different cell fractions were extracted using TRIzol™ reagent (Invitrogen, 15596026). RNA was reverse-transcribed into complementary DNA (cDNA) using the HiScript II Q RT SuperMix for qPCR (Vazyme, R223-01). RT-qPCR was conducted on a QuantStudio 3 Real-Time PCR System (Applied Biosystems) with AceQ® qPCR SYBR® Green Master Mix (Vazyme, Q121-02). The sequences of all primers used for RT-qPCR will be provided upon publication. The relative mRNA expression levels of target genes were quantified using the internal control *gapdh* for validating *BRD4* knockdown or LMNB2 for comparing transcriptional levels between rescued BRD4-WT and BRD4-12A HeLa cells, following the ΔΔCt method.

### ChIP-qPCR and ChIP-seq

Cells were crosslinked with 1% formaldehyde for 20 min at room temperature, and quenched with 0.125 M glycine. Cell pellets were rinsed twice with ice-cold PBS, lysed in lysis buffer (50 mM HEPES, pH 7.9, 5 mM MgCl_2_, 0.2% Triton X-100, 20% glycerol, 300 mM NaCl, protease inhibitor) on ice for 10 min, then centrifuged at 4 °C, 500 g for 5 min. Pellets were resuspended in SDS lysis buffer (50 mM HEPES, pH 7.4, 150 mM NaCl, 1 mM EDTA, 1% Triton X-100, 0.1% sodium deoxycholate, 0.1% SDS, protease inhibitor) and kept on ice for 30 min. Lysates were sonicated utilizing a Bioruptor sonicator (Diagenode) at high power for 30 cycles, followed by centrifugation at 17,000 g for 10 min, and supernatants collected. Antibodies were conjugated to pre-washed Protein A/G magnetic beads at 4 °C with continuous rotation for at least 2 h, followed by three times washes in SDS lysis buffer. Lysates were precleared with beads at 4 °C for 1 h with rotation, retaining 10% as input. Cleared lysates were incubated with antibody-bound beads overnight at 4 °C with rotation. Beads were washed sequentially at 4 °C with every 5 min using the following buffers: twice with SDS lysis buffer, twice with high salt wash buffer (50 mM HEPES, pH 7.5, 350 mM NaCl, 1 mM EDTA, 1% Triton X-100, 0.1% sodium deoxycholate, 0.1% SDS), twice with LiCl wash buffer (10 mM Tris-HCl, pH 8.0, 250 mM LiCl, 1 mM EDTA, 0.5% NP-40, 0.5% sodium deoxycholate) and once with TE wash buffer (10 mM Tris-HCl, pH 8.0, 1 mM EDTA, 0.2% Triton X-100), then eluted with 240 µL elution buffer (100 mM NaHCO_3_, 1% freshly added SDS) by shaking at 30 °C for 1 h. Crosslinks were reversed by adding 14.4 µL 5 M NaCl at 65 °C for 4 h, followed by adding 4 µL RNase A (Thermo Scientific, EN0531) at 37 °C for 15 min and 4 µL proteinase K (Thermo Scientific, 100005393) at 65 °C overnight. DNA was extracted by DNA clean & concentrator (Zymo Research, D4014) for ChIP-qPCR, where DNA was 1:30 diluted, and for ChIP-seq, where library preparation was performed utilizing VAHTS Universal Pro DNA Library Prep Kit (Vazyme, ND608) following the manufacturer’s protocols with 10 ng initial amount of DNA for each sample.

### ChIP-seq data analysis

ChIP-seq reads were trimmed and aligned to GRCh38/hg38 human genome using Bowtie2 (v2.4.5)^62^. Duplicates were removed with Picard (v1.88), and BAM files generated with Samtools (v1.2)^63^. Peaks were called using MACS2 (v2.2.7.1; parameters: -q 0.01 --extsize 150 --keep-dup 5)^64^. Robust BRD4 peaks (intersected from biological replicates) for both 12A and WT were merged using Bedtools (v2.31.1)^65^, and further selected when overlapping with human ENCODE blacklist regions for over 1 bp.

BRD4 binding regions were annotated as 1) active enhancers (H3K27ac^+^ H3K4me1^-^); 2) primed enhancers (H3K4me1^+^ H3K27ac^-^); 3) active promoters (H3K4me3^+^ H3K27ac^+^), and 4) primed promoters (H3K4me3^+^ H3K27ac^-^). Read counts were quantified with featureCounts (v2.0.2)^66^ and normalized to counts per million (CPM). Super-enhancers were identified by the ROSE algorithm^67^ with merged BRD4 ChIP-seq signals from BRD4-WT and BRD4-12A samples in H3K27ac regions with a stitching distance of 12.5 kb and a TSS exclusion zone of 2 kb. Coverage tracks were generated with deepTools bamCoverage with parameters ‘--normalizeUsing RPKM’ and visualized using deepTools heatmap and the Integrative Genomics Viewer (IGV)^68^.

### RNA-seq and data analysis

RNA-seq library preparation was performed utilizing VAHTS Universal V8 RNA-seq Library Prep Kit (Vazyme, NR605) following the manufacturer’s protocols with 1 µg initial amount of total RNA for each sample, and sequencing was performed on a DNBSEQ-T7 platform or a NovaSeq 6000 platform.

For RNA-seq analysis, reads were trimmed for low-quality nucleotides and adapter sequences using Trim Galore! (v0.6.7), then aligned to human genome GRCh38/hg38 with HISAT2 (v2.2.1)^69^ using the ‘-rna-strandness RF’ parameter and an additional ‘-k 30’ parameter for chromatin RNA analysis. Gene annotation GTF format files for human were downloaded from the ENSEMBL database (https://www.ensembl.org/index.html). Reads counts for genes were obtained using featureCounts (v2.0.2)^66^ and then normalized to fragments per kilobase of transcript per million mapped reads (FPKM). Differential expression analysis was performed with the R/Bioconductor package DESeq2 (v1.36.0)^70^, using *P* value < 0.05 and log_2_ |fold change| > log_2_ (1.5) as thresholds.

### DisP-seq and data analysis

DisP-seq and its analysis were performed as previously reported^47^. Briefly, isolated nuclei from 10 millions of BRD4-WT and BRD4-12A HeLa cells were resuspended in 200 μL of prewarmed MNase digestion buffer (50 mM Tris–HCl, pH 7.4, 320 mM sucrose, 4 mM MgCl_2_, 1 mM CaCl_2_) supplemented with 1× protease/phosphatase inhibitor cocktail and 0.1 mM PMSF. Chromatin digestion was initiated by adding 6 units of MNase (New England Biolabs, M0247S) and incubated at 37°C for 1 minute. The reaction was quenched by adding 800 μL ice-cold β-isox lysis buffer (20 mM Tris–HCl, pH 7.4, 187.5 mM NaCl, 5 mM MgCl_2_, 0.625% NP-40, 12.5% glycerol) supplemented with fresh 1× protease/phosphatase inhibitors, 0.1 mM PMSF, and 25 mM β-mercaptoethanol. After centrifugation of 18,400 g for 15 min at 4°C, the supernatant was collected. 10% was saved as input, and the rest was split into two parts for DMSO or 100 μM β-isox treatment. Samples were rotated at 4°C for 1 h, re-centrifuged, and supernatants discarded. Pellets were washed twice with ice-cold buffer (20 mM Tris pH 7.4, 150 mM NaCl, 5 mM MgCl_2_, 0.5% NP-40, 10% glycerol) supplemented with fresh 1× protease/phosphatase inhibitor cocktail, 0.1 mM PMSF and 20 mM β-mercaptoethanol via gentle resuspension and centrifugation of 18,400 g for 5 min at 4°C. Subsequently, 2 μL RNase (Thermo Scientific, EN0531) was added into input and pellets and incubated at 37 °C for 30 min following resuspension in 200 μL elution buffer (10 mM Tris–HCl pH 8.0, 0.1 % SDS, 150 mM NaCl and 5 mM DTT) by shaking of 600 rpm at 65 °C for 1 h. After that, proteinase K (Thermo Scientific, EO0491) were supplemented with shaking of 600 rpm for at 65 °C for 3 h, and then using DNA Clean & Concentrator (Zymo Research, D4014) to extract DNA. 2 ng DNA were used to prepare sequencing library utilizing VAHTS Universal Pro DNA Library Prep Kit (Vazyme, ND608) following the manufacturer’s protocols. Sequencing was performed at JMDNA Bio-Medical Technology Corporation on a DNBSEQ-T7 platform.

DisP-seq paired sequenced reads were aligned to GRCh38/hg38 human genome using Bowtie2 (v2.4.5)^62^ with duplicates removed using Picard tools. Peaks were called using MACS2 (2.2.7.1)^64^ with parameter: --nomodel -B --SPMR -f BAMPE --broad -p 0.01 --broad-cutoff 0.01. DisP-seq signals across identified peaks were quantified using featureCounts (v2.0.1)^66^ in a paired-end mode. Peaks with fold change greater than 1.5 (b-isox versus DMSO) were selected for downstream analyses. DISPbind (v1.0.2, https://github.com/rdong08/DISPbind)^47^ with default settings was employed to define DisP islands.

ChIP-seq datasets of 1,945 human transcription factors (TFs) from the ENCODE project were systematically analyzed. For each TF, the DisP-Enriched Disordered O-GlcNAcCluster Score was defined as the product of the overlap similarity between DisP-seq peaks and TF-bound genomic regions and the number of clustered O-GlcNAc sites within TF’s IDR. Scores greater than 1.1 were capped at 1.1.

### Gene ontology (GO) and gene set enrichment analysis (GSEA)

GO analysis of hyper peaks was performed using clusterProfiler (v4.2.2)^71^ and visualized with ggplot2. GSEA was performed using the gseGO and gseKEGG function in the R package with a *P* value threshold of 0.05. Enriched pathways were visualized using the gseaNb function from the GseaVis R package, with significant pathways highlighted.

Target members were analyzed using the STRING database (version 12.0) with the following settings: species as *Homo sapiens*, interaction score as 0.9, and the option to hide disconnected nodes. The output results were then imported into Cytoscape v3.7.2, and the top three GO terms were selected to display the context of protein-protein interactions (PPIs).

### Statistics and reproducibility

The immunoblotting and immunofluorescence data were quantified using Fiji ImageJ. Data analysis and statistical diagrams were conducted using GraphPad Prism 8 or R software version 4.2.3. Unpaired two-tailed t-tests were used for comparisons between two groups, and two-way ANOVA tests were used for comparisons among data sets in the *in vitro* droplet formation assay. A *P* value less than 0.05 was considered significant. Exact *P* values were reported in each figure panel, while very small *P* values were shown as a ’less-than’ range. The Pearson correlation coefficient (PCC) or Spearman’s rank correlation coefficients (Spearman coeff) was calculated to evaluate correlation, and R with *P* value in scatter plots indicates the proportion of variance in the dependent variable explained by a linear relationship. In box plots, the center line represents the median, box limits indicate the upper and lower quartiles, and *P* values were calculated using the non-parametric Wilcoxon-Mann-Whitney test (Wilcoxon rank sum test, two-sided).

### Data availability and public datasets

The mass spectrometry data of BRD4 glycosites and chromatin-bound proteome have been deposited to the ProteomeXchange Consortium through the iProX^72,73^ repository with the dataset identifier PXD057479, which will be released to public upon publication. The sequence data generated in this study have been deposited in the NCBI Gene Expression Omnibus (GEO) under the accession number GSE282031. The public ChIP-seq datasets of were downloaded from GEO database: H3K27ac (GSM733684), H3K4me1 (GSM798322), H3K4me3 (GSM733682) ChIP-seq data in HeLa cells, as well as YTHDC1 ChIP-seq data in MCF7 cellls (GSE143441). The unpublished HeLa chromatin-associated meRIP-seq data generated by Jun Liu’s lab was used for functional analysis of RNA methylation.

---

**Extended Data Fig. 1.**
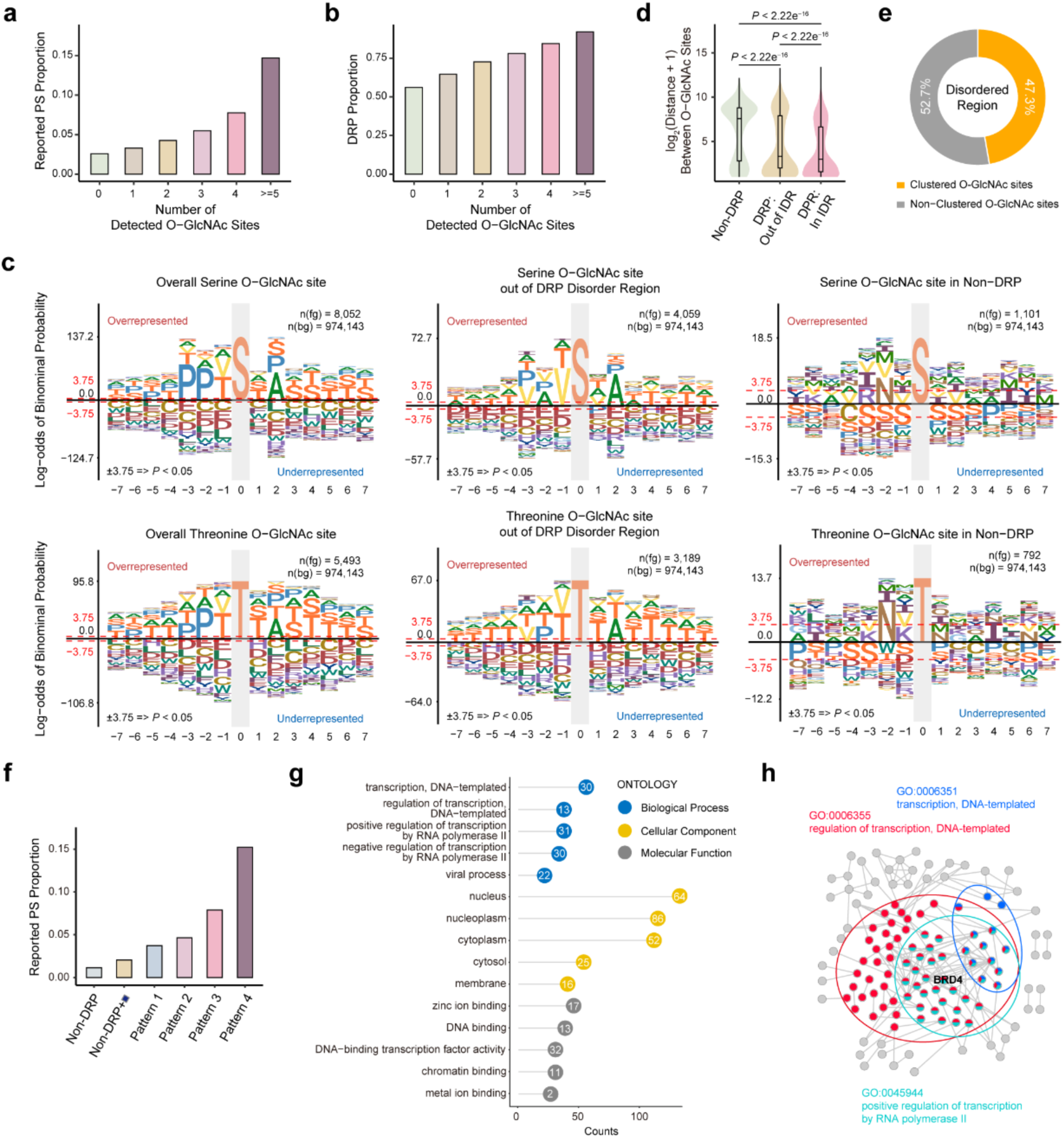
Phase-separating proteins with intrinsically disordered regions (IDRs) tend to enrich O-GlcNAc clusters. **a.** Bar chart showing the proportion of reported phase-separating (PS) proteins across proteins with different numbers of O-GlcNAc sites. **b.** Bar chart showing the IDR-containing proteins (DRP) proportion across proteins with different numbers of O-GlcNAc sites. **c.** O-GlcNAcylation consensus sequences of the entire human O-GlcNAcome (left), of glycosites outside IDRs of DRPs (middle), and of glycosites in non-DRPs. **d.** Box-violin plot showing the amino acid distance between O-GlcNAc sites within different regions. Two-sided Wilcoxon rank-sum tests were used for statistical analyses. **e.** Pie chart showing the proportion of clustered and non-clustered O-GlcNAc sites within disordered regions. **f.** Bar chart showing the reported PS proportion across different O-GlcNAc site distribution patterns shown in **Fig.1k**. **g.** Gene Ontology (GO) analysis of predicted PS proteins containing O-GlcNAc clusters. GO terms associated with molecular function, cellular component, and biological processes are highlighted in gray, yellow, and blue circles with -log_10_ *P* inside, respectively. **h.** Protein-protein interaction (PPI) network of proteins analyzed in **g**, with proteins associated with three GO terms highlighted in green, red and blue.

**Extended Data Fig. 2.**
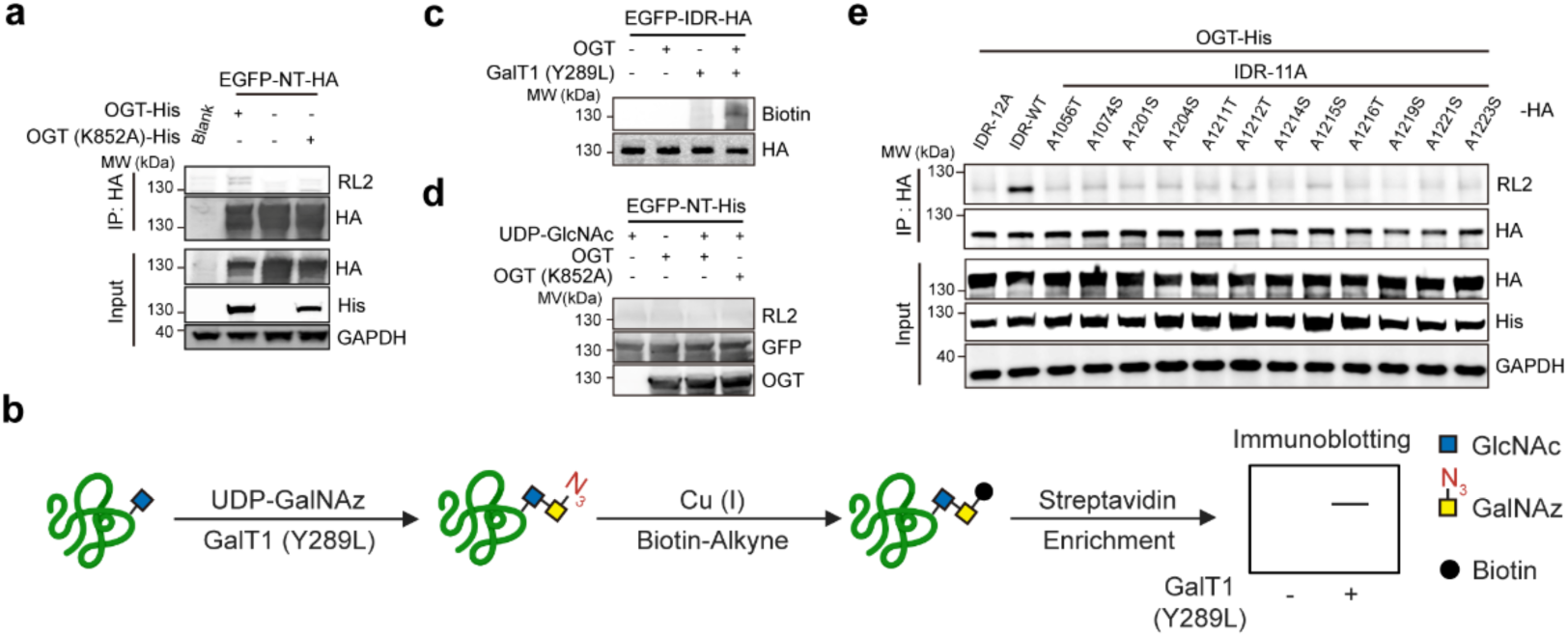
O-GlcNAcylation validation on indicated BRD4 variants. **a.** Immunoblots showing O-GlcNAc level on HA and EGFP-tagged BRD4-NT (EGFP-NT-HA) when co-expressed with OGT or OGT (K852A) in HEK293T cells. **b.** Scheme of the chemoenzymatic labeling process followed by click chemistry for O-GlcNAcylated protein enrichment. **c.** Immunoblots showing the validation of O-GlcNAc modification on HA-tagged BRD4-IDR (EGFP-IDR-HA) when co-expressed with OGT in HEK-293T cells following the chemoenzymatic labeling assay shown in **b**. **d.** Immunoblots showing O-GlcNAc level on purified EGFP-fused BRD4-NT-His (EGFP-NT-His) after *in vitro* O-GlcNAcylation assay. **e.** Immunoblots showing O-GlcNAc levels on BRD4-IDR, BRD4-IDR-12A, and BRD4-IDR mutants preserving a single O-GlcNAc site, when co-expressed individually with OGT-His in HEK293T cells. RL2, anti-O-GlcNAc antibody (RL2). Data in **a, c–e** represent at least two biological replicates.

**Extended Data Fig. 3.**
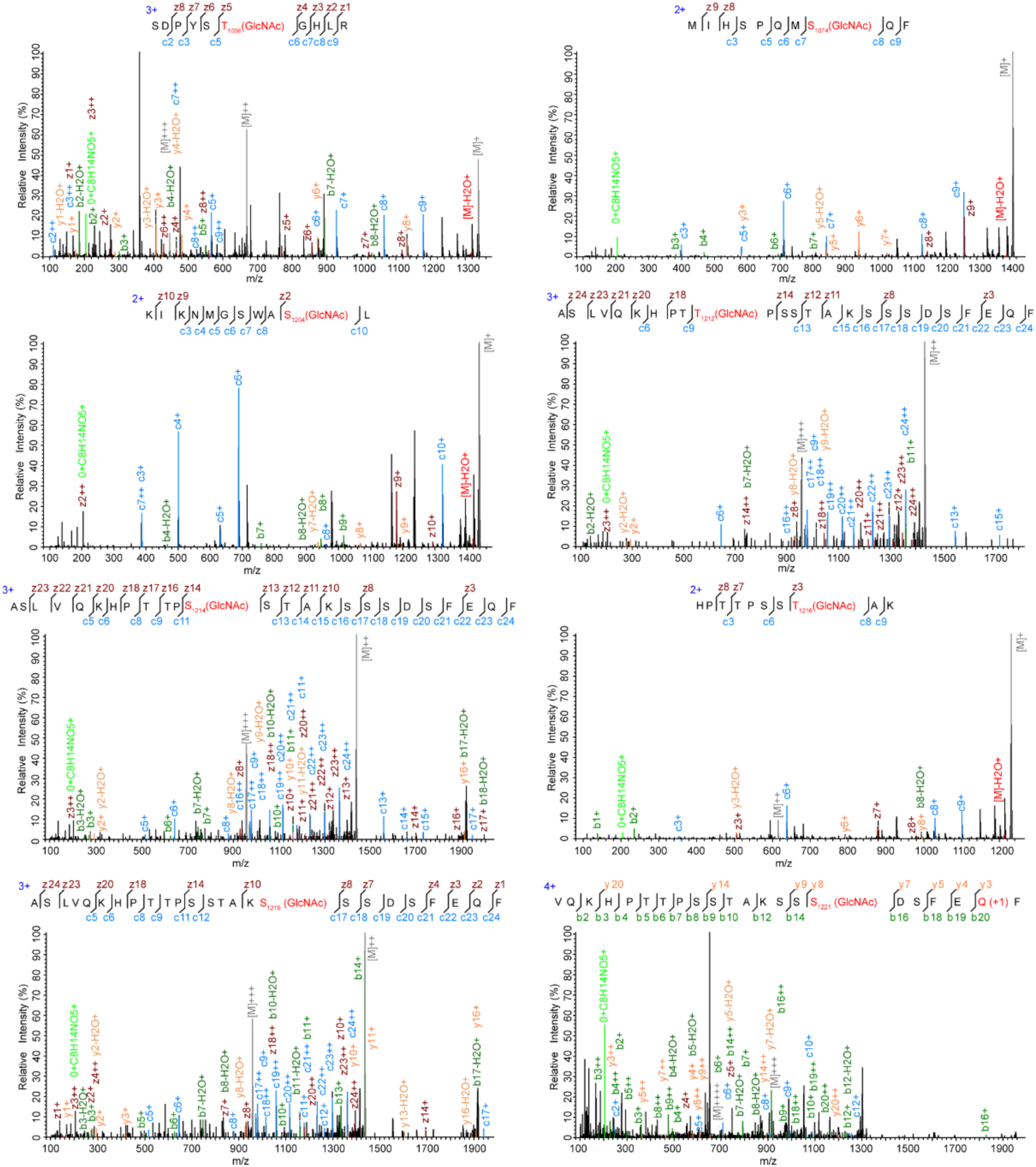
Representative mass spectra of BRD4-IDR peptides with T1056, S1074, S1204, T1212, S1214, T1216, S1219, and S1221 sites modified with O-GlcNAc. HexNAc, b, y, and c, z ions were annotated.

**Extended Data Fig. 4.**
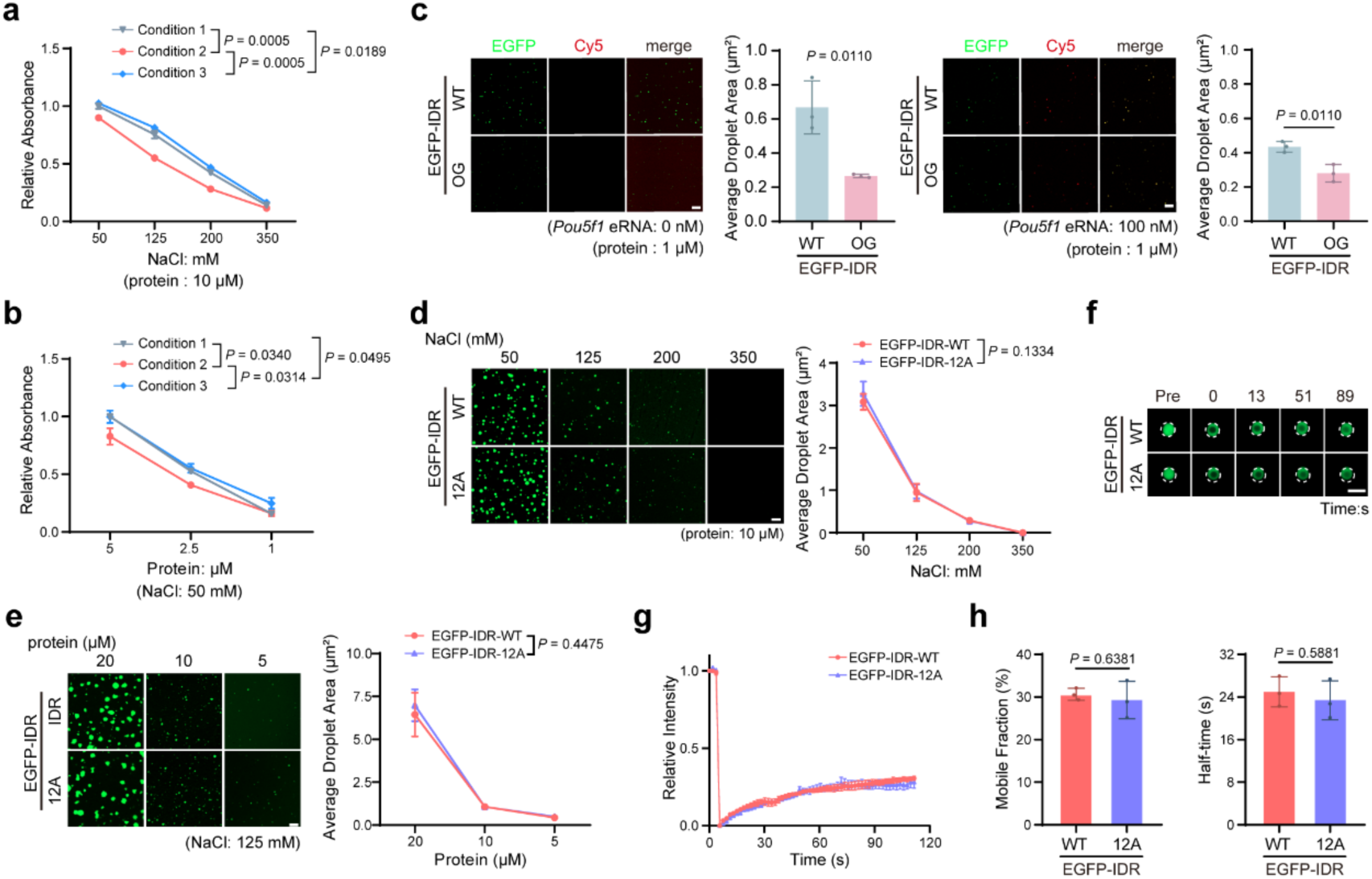
O-GlcNAcylation reduces BRD4-IDR condensates *in vitro*, whereas the 12A mutation causes negligible effects. **a, b.** Quantitative results of EGFP-BRD4-IDR droplet formation under different conditions at different NaCl (**a**) or protein concentrations (**b**) using turbidity assays. Three conditions are the same as conditions shown in **Fig. 3a, c**. **c.** Representative images and quantification results of the droplet formation assay using 1 μM EGFP-BRD4-IDR (EGFP-IDR) or its O-GlcNAc-modified form (EGFP-IDR-OG) droplets, mixed with the indicated concentrations of *Pou5f1* enhancer RNA (indicated in the Cy5 channel). **d.** Representative images (left) and quantification (right) of droplet formation using 10 μM EGFP-IDR or its mutant EGFP-IDR-12A at different NaCl concentrations. **e.** Representative images (left) and quantification (right) of droplet formation using EGFP-IDR or EGFP-IDR-12A with different protein concentrations at 125 mM NaCl. **f, g.** Representative FRAP images (**f**) and curves (**g**) of EGFP-IDR and EGFP-IDR-12A. **h.** Histograms showing the mobile fraction and recovery half-time of droplets in **f** after FRAP. Scale bar, 5 μm (**f**), 10 μm (**c, d, e**). Data are the mean ± s.d. from n = 3 biologically independent samples. Statistical analyses were performed using two-way ANOVA tests in **a, b, d, e**, and two-tailed t-tests in **c, h**.

**Extended Data Fig. 5.**
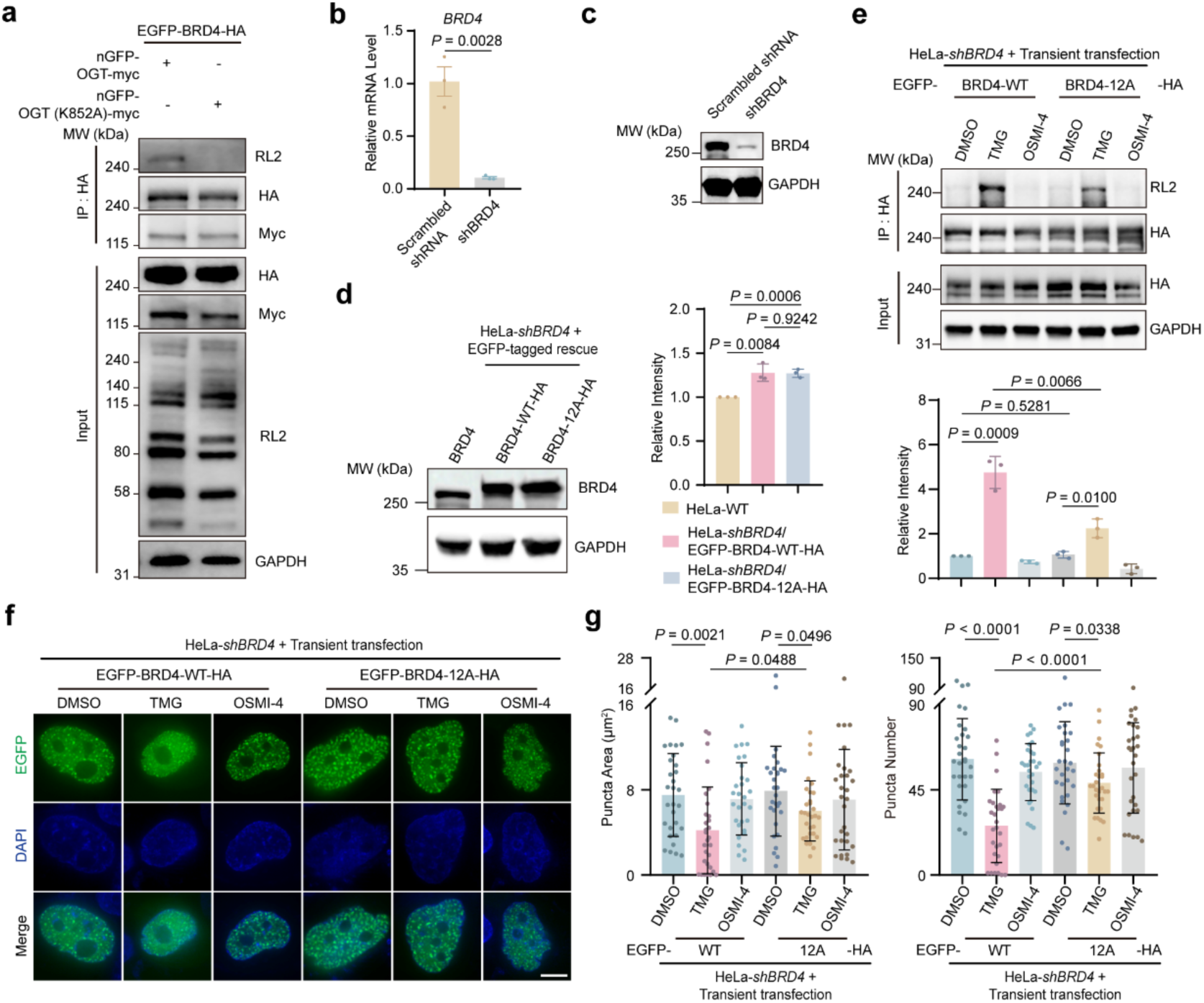
O-GlcNAcylation diminishes BRD4 condensates in HeLa cells. **a.** Immunoblots showing O-GlcNAc level of EGFP-BRD4-HA after targeted glycosylation using mCherry-tagged nGFP-OGT or nGFP-OGT (K852A) in HeLa cells. RL2, anti-O-GlcNAc antibody (RL2). **b.** Histogram showing quantitative real-time PCR (qRT-PCR) results of *BRD4* mRNA levels in scrambled *shRNA* and *shBRD4* knockdown cells. **c.** Immunoblots showing BRD4 protein level after shRNA knockdown in HeLa cells. **d.** Immunoblots (left) and quantification (right) of BRD4 protein levels in *shBRD4* knockdown HeLa cells rescued with EGFP-tagged BRD4-WT-HA or BRD4-12A-HA. **e.** Representative immunoblots (top) and quantification (bottom) of O-GlcNAc levels of EGFP- and HA-tagged BRD4-WT or BRD4-12A transiently expressed in *shBRD4* HeLa cells after the treatments of DMSO, TMG or OSMI-4 for 48 h. **f.** Live-cell images of puncta formed by EGFP- and HA-tagged BRD4-WT or BRD4-12A when transiently expressed in *shBRD4* HeLa cells under the treatments of DMSO, TMG or OSMI-4 for 48 h. Scale bar, 5 μm. **g.** Quantified puncta areas (left) and numbers (right) in samples in **f**. n = 30 cells per group, pooled from three independent replicates. Data in **a, c** represent at least two biological replicates. n = 3 biologically independent experiments in **d, e**. Data in **b, d, e, g** are the mean ± s.d.. Two-tailed t-tests were used for statistical analyses.

**Extended Data Fig. 6.**
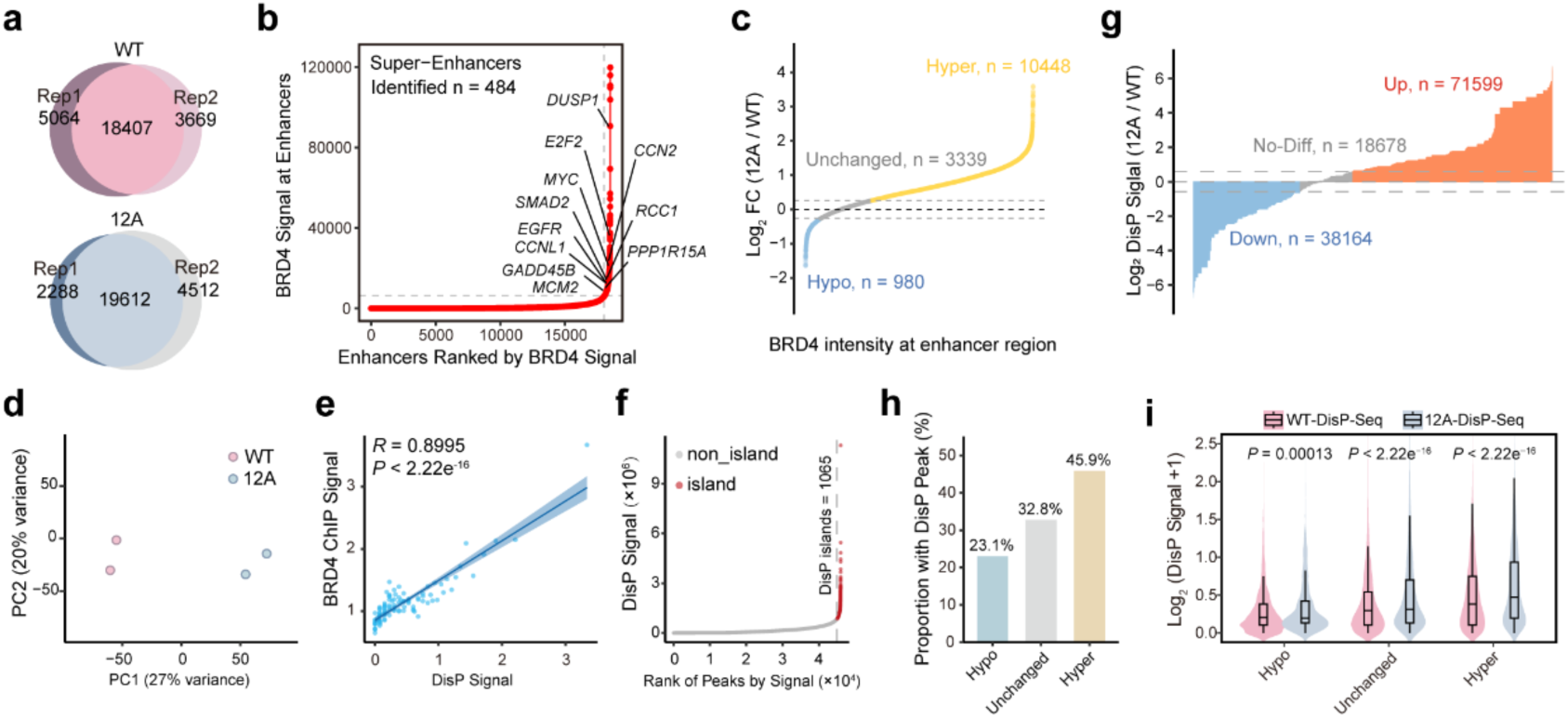
BRD4 12A exhibits higher occupancy and LLPS-related signal at enhancer regions. **a.** Venn diagrams showing the consistency between two replicates of ChIP-seq peaks from BRD4-WT and BRD4-12A cells. **b.** Rank ordered plots of enhancers ranked by BRD4 ChIP-seq signal. The dotted line indicates the super-enhancer (SE) threshold level set by the ROSE algorithm, with a total of 484 SEs. **c.** Distribution plot showing ChIP-seq signal fold changes of BRD4-12A versus BRD4-WT cells at enhancers. FC, fold change. **d.** Principal component analysis (PCA) score plot showing two replicates of DisP-seq in BRD4-WT and BRD4-12A cells. **e.** The correlation between BRD4 ChIP-seq signal and DisP-seq signal. Peaks were categorized into 100 bins based on the intensity rank of DisP Signal. The Pearson correlation coefficient was used to determine the correlation. **f.** Identification of DisP islands according to DisP-seq signal. **g,** Distribution plot showing DisP-seq signal fold changes of BRD4-12A versus BRD4-WT cells. No-Diff, no-difference. **h.** Bar chart showing the proportion of DisP-seq signals detected across three ChIP-seq categories (Hypo, Unchanged, and Hyper peaks in BRD4-12A versus BRD4-WT cells). **i.** Box-violin plot showing DisP-seq signal across three ChIP-seq categories (Hypo, Unchanged, Hyper peaks in BRD4-12A versus BRD4-WT cells). Two-sided Wilcoxon rank-sum tests were used for statistical analyses.

**Extended Data Fig. 7.**
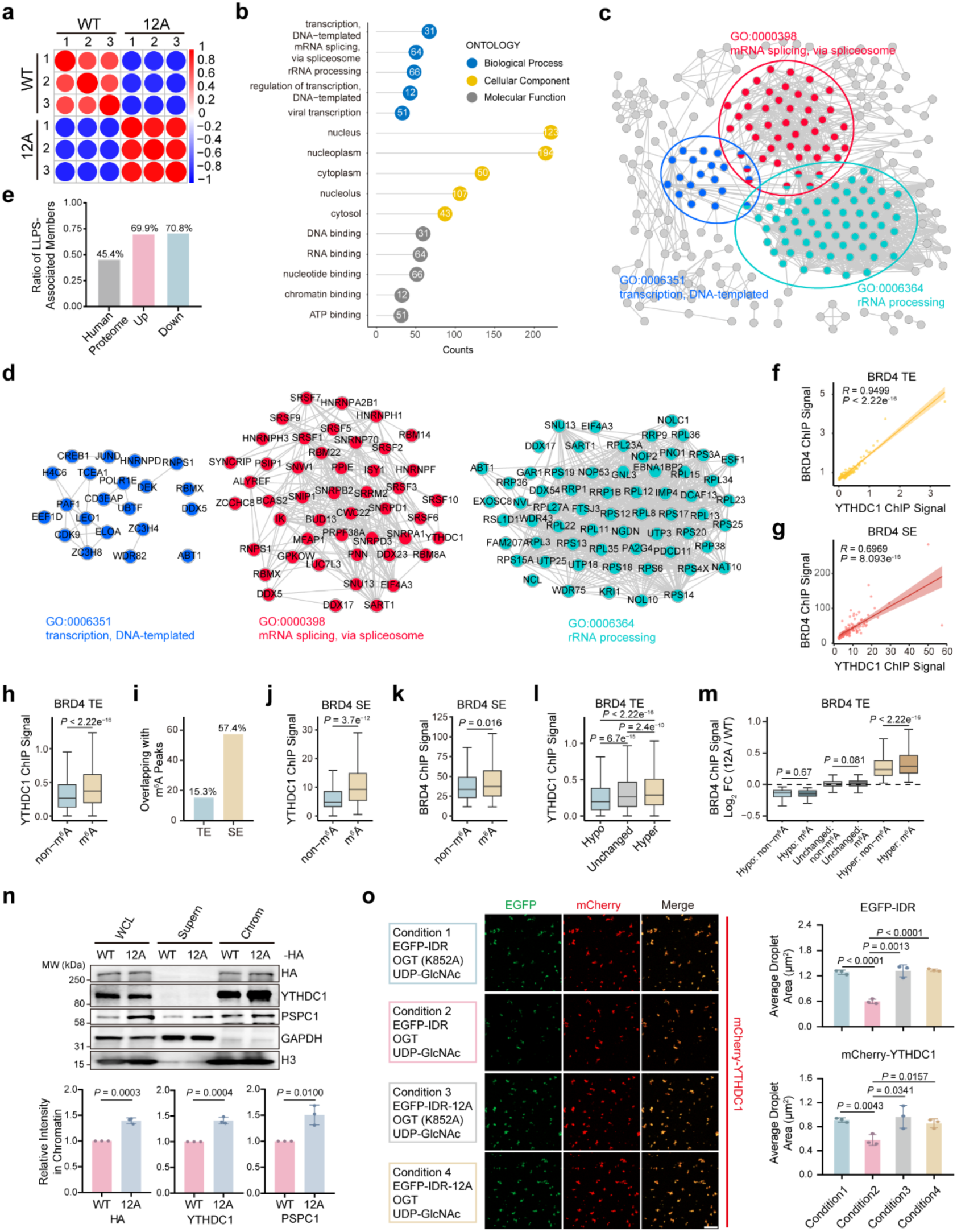
YTHDC1 facilitates the recruitment of BRD4 to the chromatin. **a.** Hierarchical clustered heatmap showing pairwise chromatin-MS correlations between BRD4-WT and BRD4-12A cells. Pearson’s correlation was used to determine coefficients. **b.** Gene Ontology (GO) analysis of upregulated proteins identified by chromatin-MS in BRD4-12A versus BRD4-WT cells. GO terms associated with molecular function, cellular component, and biological processes are highlighted in gray, yellow, and blue circles with -log_10_ *P* inside, respectively. **c.** Protein-protein interaction (PPI) network of upregulated proteins analyzed in **b**. **d.** Proteins associated with three GO terms (“transcription, DNA-templated”, “mRNA splicing, via spliceosome”, or “rRNA processing”) are annotated and highlighted in blue, red and green. **f, g.** Correlations between BRD4 ChIP-seq signal and YTHDC1 ChIP-seq signal at BRD4-bound TEs (**f**) and SEs (**g**). Peaks were categorized into 100 bins based on the intensity rank of YTHDC1 Signal. Pearson’s correlation coefficients (R) with *P* values were used in **f, g. h.** Boxplot showing YTHDC1 ChIP-seq signal in m^6^A- or non-m^6^A-modified regions at BRD4 TEs. **i.** Bar chart showing the proportion of m^6^A modified peaks in TE and SE regions. **j.** Boxplot showing YTHDC1 ChIP-seq signal in m^6^A- or non-m^6^A-modified regions at SEs. **k.** Boxplot showing BRD4 ChIP-seq signal in m^6^A- or non-m^6^A-modified regions at SEs. **l.** Boxplot showing YTHDC1 ChIP-seq signal across three BRD4 ChIP-seq categories at TEs (Hypo, Unchanged, and Hyper peaks in BRD4-12A versus BRD4-WT cells). **m.** Boxplot showing fold changes of BRD4 ChIP-seq signal in BRD4-12A versus BRD4-WT cells in m^6^A- and non-m^6^A-modified sites across three BRD4 ChIP-seq categories at TEs. FC, fold change. **n.** Immunoblots and quantification of chromatin-bound BRD4, PSPC1 and YTHDC1 in BRD4-12A and BRD4-WT cells. **o.** Representative images (left) and quantified (right) droplets formed by 2 μM mCherry-YTHDC1 mixed with 5 μM EGFP-IDR-WT or EGFP-IDR-12A under indicated conditions. Scale bar, 10 μm. Data in **n, o** are the mean ± s.d.. Two-tailed t-tests were used in **n, o** and two-sided Wilcoxon rank-sum tests were used in **h–m** for statistical analyses.

**Extended Data Fig. 8.**
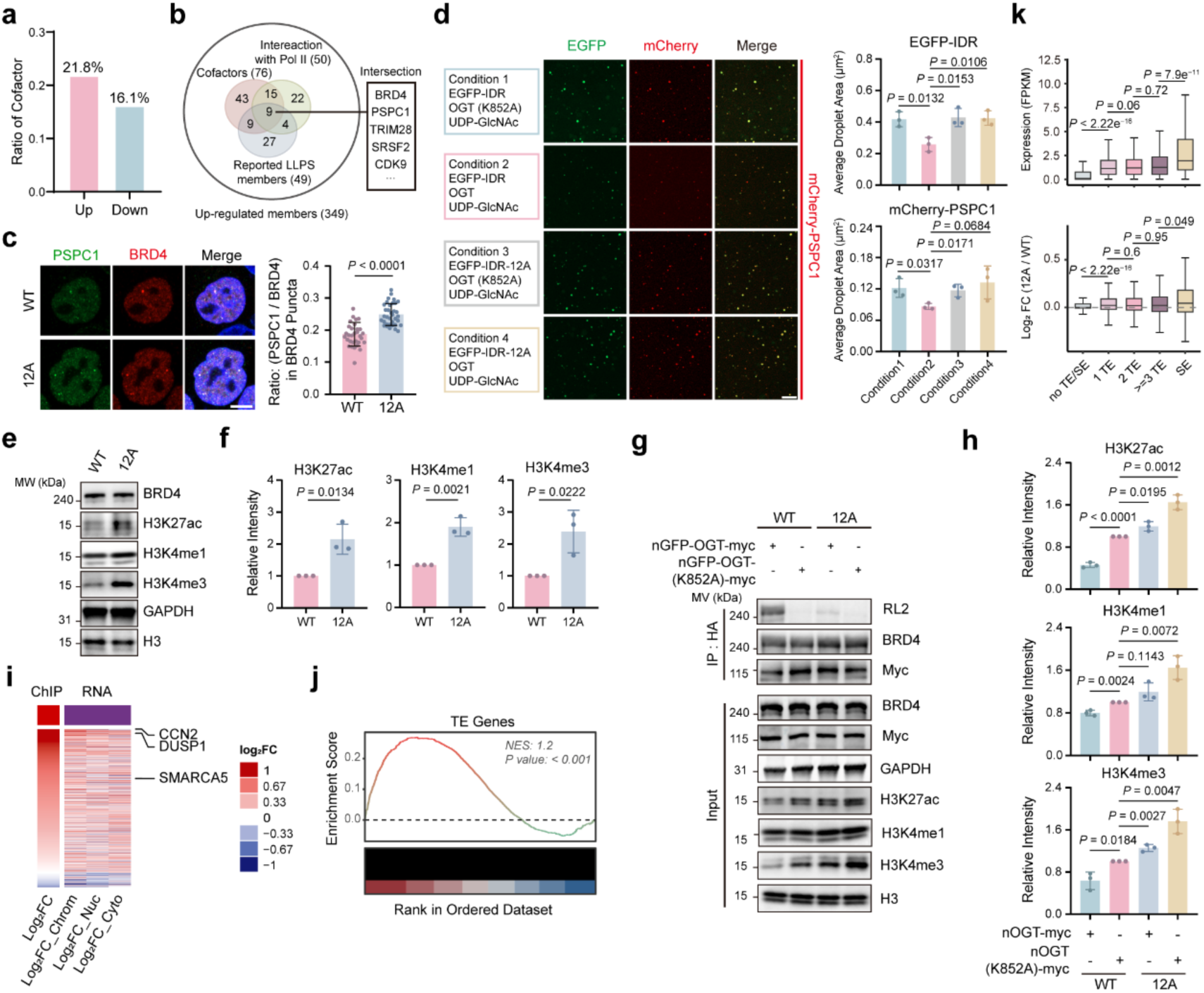
Loss of O-GlcNAc on BRD4 promotes transcription. **a.** Bar chart showing the ratio of transcription cofactors in upregulated and downregulated proteins identified by the chromatin-MS shown in **Fig. 5m**. **b.** Venn diagram showing the overlapped nine proteins by intersecting upregulated proteins, reported LLPS members, transcription cofactors, and RNA Pol II-interacting proteins. Representative proteins are listed in the right. **c.** Representative immunofluorescence images and quantified results showing the localization and fluorescence density ratio of PSPC1 and BRD4 in BRD4 puncta at WT and 12A cells. n = 35 and 32 cells in WT and 12A cells, respectively, pooled from three independent replicates. **d.** Representative images (left) and quantified (right) droplets formed by 5 μM mCherry-PSPC1 mixed with 5 μM EGFP-IDR-WT or EGFP-IDR-12A under indicated conditions. **e, f,** Immunoblots (**e**) and quantification (**f**) of modification levels of three histone markers H3K27ac, H3K4me1 and H3K4me3 in BRD4-WT and BRD4-12A cells. **g, h.** Immunoblots (**g**) and quantification (**h**) of modification levels of three histone markers H3K27ac, H3K4me1 and H3K4me3 in BRD4-WT and BRD4-12A cells undergoing targeted glycosylation using nGFP-OGT. RL2, anti-O-GlcNAc antibody (RL2). **i.** Heatmap showing comparisons of log_2_ FC among BRD4 ChIP-seq signals and RNA-seq results from indicated cell fractions in BRD4-12A versus BRD4-WT cells. Chrom, chromatin; Nuc, nucleoplasm; Cyto, cytoplasm. **j.** Gene set enrichment analysis (GSEA) plots showing expression differences of hyper genes in BRD4-12A versus BRD4-WT ChIP-seq results. NES, normalized enrichment score. **k.** Boxplots showing gene expression (top) and FCs (bottom) between 12A versus WT cells across different TE numbers as well as SE. Scale bar, 5 μm (**c**), 10 μm (**d**). Two-sided Wilcoxon rank-sum tests were used for statistical analyses. FC, fold change. Data in **c, d, f, h** are the mean ± s.d. and analyzed using two-tailed t-tests.

**Extended Data Fig. 9.**
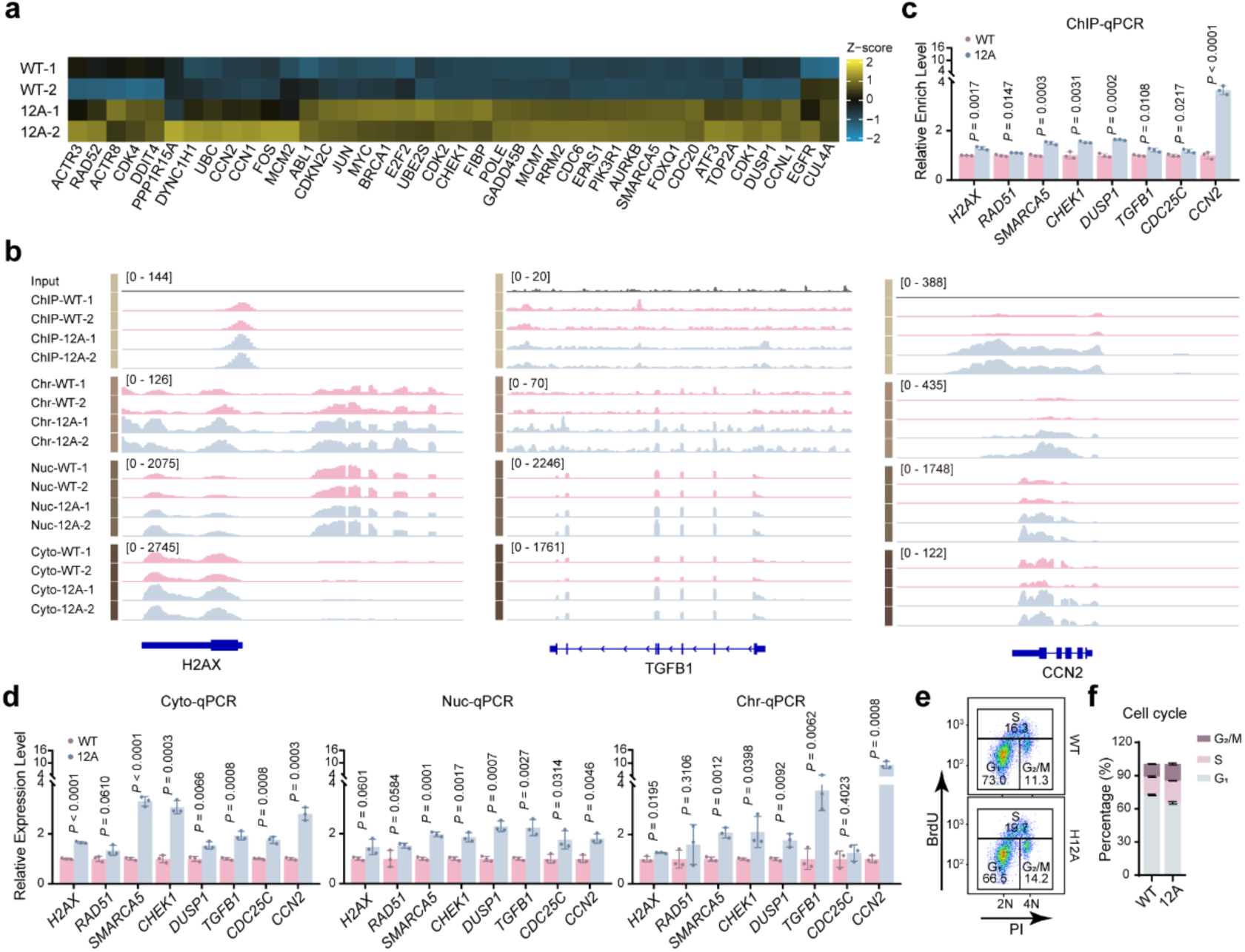
O-GlcNAcylation on BRD4 inhibits expression of genes related to DNA damage repair, cell cycle, and proliferation. **a.** Heatmap showing representative gene expression levels associated with key biological processes in BRD4-12A and WT cells. **b.** Integrative Genomics Viewer plots showing BRD4 ChIP-seq binding signals and RNA-seq expression levels of H2AX, TGFB1, and CCN2 in BRD4-WT or 12A cell fractions. Chr, chromatin; Nuc, nucleoplasm; Cyto, cytoplasm. **c, d.** Histograms showing BRD4 ChIP-quantitative PCR (ChIP-qPCR, **c**) and qRT-PCR (**d**) of indicated genes in 12A and WT cells. Genes were selected from upregulated genes in **Fig. 6e, f**. **e, f.** Flow cytometric cell cycle analysis (**e**) and quantification (**f**) of BRD4-12A and BRD4-WT cells without synchronization. n = 3 biologically independent experiments. Data in **c, d, f** represent the mean ± s.d. from n = 3 independent experiments. Two-tailed t-tests were used for statistical analyses.

